# Temporal proximity proteomics reveals ANP32A roles in replication fork progression and end joining

**DOI:** 10.64898/2026.09.11.750924

**Authors:** Carla-Marie Jurkovic, Jennifer Raisch, Gwendoline Marbach, Dominique Lévesque, Isabelle Marois, Billel Djerir, Alberto David Delgado Monterroso, Alexandre Maréchal, François-Michel Boisvert

**Affiliations:** Department of Immunology and Cell Biology, Faculty of Medicine and Health Sciences, Université de Sherbrooke; Department of Biology, Faculty of Science, Université de Sherbrooke

**Keywords:** ANP32A, Replication stress, Proximity proteomics, AirID, Chromatin, DNA replication, Replication fork progression, DNA damage response, Non-homologous end joining, Genome maintenance

## Abstract

Acidic nuclear phosphoprotein 32 family member A (ANP32A) regulates chromatin and histone homeostasis, but its contribution to DNA replication and repair remains unclear. Time-resolved AirID proximity labeling with data-independent acquisition mass spectrometry revealed phased remodeling of the ANP32A proximal proteome during hydroxyurea-induced replication stress. Early stress enriched DNA replication and repair factors, including FEN1, XRCC5/Ku80, and XRCC6/Ku70, whereas prolonged exposure shifted the network towards checkpoint and cell-cycle regulators. Orthogonal proximity ligation assays revealed sustained ANP32A-FEN1 proximity and transiently increased ANP32A-XRCC5 proximity. ANP32A was dynamically recruited to γH2AX-marked lesions following laser microirradiation and showed increased proximity to γH2AX after hydroxyurea treatment. ANP32A loss impaired proliferation and G1/S progression and moderately reduced replication fork velocity. However, it did not promote nascent strand degradation at stalled forks or measurably alter homologous recombination reporter activity. In contrast, ANP32A loss reduced non-homologous end-joining reporter activity, consistent with its stress-induced proximity to Ku proteins. These findings identify ANP32A as a chromatin-associated factor supporting replication fork progression and end joining while dispensable for stalled-fork protection, linking ANP32A-dependent chromatin regulation to genome maintenance.

**HIGHLIGHTS:**

- Replication stress dynamically remodels the ANP32A proximal proteome
- ANP32A loss impairs G1/S progression and slows replication forks
- ANP32A is recruited to γH2AX-marked DNA lesions during replication stress
- ANP32A promotes end joining but is dispensable for stalled fork protection

**IN BRIEF:** Jurkovic et al. show that replication stress dynamically remodels the ANP32A proximal proteome. ANP32A supports G1/S progression, replication fork progression, and end joining but is dispensable for stalled fork protection, linking ANP32A-dependent chromatin regulation to genome maintenance.

## INTRODUCTION

Faithful duplication of the genome requires replication forks to progress through a complex chromatin landscape while continuously encountering obstacles to DNA synthesis. DNA lesions, difficult-to-replicate sequences, transcription-replication conflicts, and nucleotide depletion can slow or stall fork progression, collectively generating replication stress (Muñoz and Méndez 2016). Although stalled forks can often be stabilized and restarted, failure to protect or properly process them can result in fork degradation or collapse, leading to DNA breaks, chromosome rearrangements, and genomic instability (Macheret and Halazonetis 2015; Zeman and Cimprich 2015; Krenning et al. 2019). Cells therefore mount a coordinated response that integrates checkpoint signaling, fork protection and remodeling, replication restart, and DNA repair. This response requires rapid and dynamic reorganization of the protein and chromatin environment surrounding stalled forks (Liao et al. 2018; Saldanha et al. 2023). Proteomic approaches that capture nascent DNA or proteins in the immediate vicinity of the replisome have begun to define this reorganization with temporal resolution (Dungrawala et al. 2015; Jurkovic et al. 2024). These studies show that fork stalling triggers extensive, time-dependent changes in protein association, encompassing canonical replication, checkpoint, and DNA repair machinery, as well as chromatin regulators and multifunctional nuclear proteins. Yet proximity to stalled forks alone does not distinguish proteins that act directly in fork remodeling, protection, or restart from those that modify the surrounding chromatin or participate in downstream signaling and repair. Consequently, the mechanistic contributions of many stress-regulated fork-associated proteins remain unresolved. Defining these noncanonical components is essential to understand how replication, chromatin organization, and DNA repair are coordinated to preserve genome stability.

One such factor is ANP32A, a member of the acidic nuclear phosphoprotein 32 kDa (ANP32) family. The core mammalian ANP32 paralogs, ANP32A, ANP32B, and ANP32E, are small and evolutionarily conserved proteins defined by an N-terminal leucine-rich repeat domain and a C-terminal low-complexity acidic region (Chen et al. 2008; Reilly et al. 2014; Ivovič et al. 2023). Despite this shared architecture, ANP32 proteins participate in diverse cellular processes, including transcriptional control, histone regulation, nucleocytoplasmic transport, apoptosis, and viral genome replication (Seo et al. 2002; Jiang et al. 2003; Staller et al. 2019; Wang et al. 2019; Long et al. 2019). Their functions can be overlapping in some contexts yet highly paralog-specific in others, as illustrated by the partially redundant requirements for ANP32A and ANP32B during influenza replication and the selective activity of ANP32E as an H2A.Z histone chaperone (Obri et al. 2014; Staller et al. 2019). Thus, despite extensive functional annotations, the molecular basis of ANP32 paralog specialization and their contributions to mammalian genome maintenance remain poorly defined.

ANP32A is particularly intriguing in this context because several observations connect it to genome duplication and replication stress. ANP32A/PP32 associates with newly synthesized histone H4 and restrains premature HAT1-dependent acetylation; depletion of ANP32A disrupts H4 maturation and causes S-phase accumulation (Saavedra et al. 2017). Time-resolved iPOND analyses have also detected ANP32A at nascent DNA, with its recovery varying across HU exposure and ATR-inhibited conditions (Dungrawala et al. 2015). In our previous mapping of 17 replisome-proximal interactomes, ANP32A was similarly identified as an HU-regulated proximity partner of multiple replication and histone-chaperone proteins (Jurkovic et al. 2024). Moreover, genome-scale CRISPR screens revealed condition-dependent fitness effects of ANP32A loss following chronic or acute HU exposure and ionizing radiation (Olivieri et al. 2021). Together, these findings place ANP32A at the intersection of chromatin assembly, stalled-fork responses, and genome maintenance, but do not establish whether it acts directly at replication forks or instead coordinates downstream cell-cycle and DNA repair pathways.

Here, we used AirID-based proximity labeling coupled with quantitative mass spectrometry to define the ANP32A proximal proteome under basal conditions and throughout HU-induced replication stress. This time-resolved analysis revealed dynamic associations with proteins involved in DNA replication, cell-cycle regulation, and DNA repair, including FEN1 and the non-homologous end-joining (NHEJ) factors XRCC5 and XRCC6. Functional studies showed that ANP32A loss impairs G1/S progression, slows replication-fork progression, and selectively reduces NHEJ efficiency, while leaving stalled fork protection and homologous recombination largely intact. Together, these findings identify ANP32A as a stress-responsive, chromatin-associated factor that supports replication-fork progression and NHEJ, thereby expanding the functions of the ANP32 family in mammalian genome maintenance.

## METHODS

### RESOURCE AVAILABILITY

#### Lead contact

Further information and requests for resources and reagents should be directed to and will be fulfilled by the lead contact, François-Michel Boisvert.

#### Materials availability

The AirID-3×Myc-ANP32A construct and the ANP32A-knockout cell lines generated in this study are available from the lead contact upon reasonable request.

#### Data and code availability

The mass spectrometry proteomics data have been deposited with the ProteomeXchange Consortium through the PRIDE repository (Perez-Riverol et al. 2022) with the dataset identifier PXD082823. All other data supporting the conclusions of this study are included in the article and its supplemental information. Any additional information required to reanalyze the data is available from the lead contact upon reasonable request.

## EXPERIMENTAL MODELS

### Cell lines and culture conditions

Human U2OS osteosarcoma cells (ATCC, HTB-96) and U2OS Flp-In T-REx (U2OSFT) cells (Thermo Fisher Scientific) were maintained in high-glucose Dulbecco’s modified Eagle’s medium containing 4.5 g/L glucose, L-glutamine, and sodium pyruvate (Wisent, 319-005-CL). The medium was supplemented with 10% fetal bovine serum (VWR, CA76322-116), 20 IU/mL penicillin, 20 μg/mL streptomycin, and 10 mM HEPES (Wisent, 450-201-EL and 330-050-EL). Cells were maintained at 37°C in a humidified atmosphere containing 5% CO₂. U2OSFT cells were maintained under hygromycin B and blasticidin selection as described below. All cell lines were tested for mycoplasma.

## METHOD DETAILS

### Generation of inducible AirID cell lines

The ANP32A coding sequence was amplified from U2OS cDNA using TransStart KD Plus DNA polymerase (TransGen Biotech, AP301-01) and primers containing attB recombination sites. The PCR product was transferred into the pDONR221 entry vector using Gateway BP Clonase II (Thermo Fisher Scientific, 11789020). Following transformation into DH10β competent bacteria and selection with kanamycin, the resulting entry clone was verified by Sanger sequencing.

ANP32A was subsequently transferred by Gateway LR recombination into a modified pgLAP1 expression vector encoding an N-terminal AirID-3×Myc tag. An AirID-3×Myc-construct lacking ANP32A but supplemented with NLS sequence was generated for use as a proximity-labeling control. LR reaction products were transformed into DH10β bacteria, selected with ampicillin, and verified by sequencing.

U2OSFT cells were cotransfected with the AirID-3×Myc-ANP32A or AirID-3×Myc expression vector and pOG44 Flp recombinase using Lipofectamine 2000. Stable integrants were selected in medium containing 100 μg/mL hygromycin B (Wisent, 450-141-XL) and 100 μg/mL blasticidin-HCl (Wisent, 450-190-XL). Unless otherwise indicated, expression of the AirID constructs was induced with 2.5 μg/mL doxycycline for 48 h.

### Generation of ANP32A-knockout cells

ANP32A-knockout U2OS cells were generated using CRISPR-Cas9 genome editing based on the protocol described by (Ran et al. 2013). A single-guide RNA targeting ANP32A (5’-CACCGTTCTTAAGTACAATCAACGT-3’) was selected using the Invitrogen CRISPR design tool and cloned into pSpCas9(BB)-2A-GFP (PX458; Addgene, 48138). The resulting plasmid was verified by Sanger sequencing.

U2OS cells were cotransfected with the ANP32A-targeting PX458 construct and pEGFP-Puro (Addgene, 45561) using jetPRIME transfection reagent (VWR, CA89129-924). Transfected cells were selected with 2.5 μg/mL puromycin (Wisent, 400-160-EM) for 5 days. Single-cell-derived colonies were isolated, expanded, and screened by PCR amplification of the targeted genomic region. Editing was evaluated using the T7 endonuclease I assay and Sanger sequencing followed by TIDE analysis (Brinkman et al. 2014). Complete loss of ANP32A protein was confirmed by immunoblotting. Two independently derived knockout clones, ANP32A KO 3.1 and KO 3.2, were used throughout the study.

### Cell proliferation assay by crystal violet staining

U2OS WT, ANP32A KO 3.1, and ANP32A KO 3.2 cells were seeded at 8,000 cells per well in separate 12-well plates designated for collection at 0, 24, 48, 72, 96, or 120 h. Cells were cultured without medium replacement. The 0 h (hour) plate was fixed on the morning after seeding, after cell attachment but before substantial proliferation. At each time point, cells were washed with PBS and fixed with 1% glutaraldehyde in PBS for 5 min at room temperature with gentle agitation. Cells were washed twice with PBS and stored in PBS at 4°C until all time points had been collected.

Cells were stained with 0.1% crystal violet in PBS for 30 min at room temperature, washed by gentle immersion in running water, and air-dried. Plates were imaged before the bound dye was solubilized in 10% acetic acid with agitation. Absorbance was measured at 590 nm and normalized to the corresponding 0 h measurement to calculate relative cell growth.

### Cell-cycle analysis by propidium-iodide staining

U2OS WT, ANP32A KO 3.1, and ANP32A KO 3.2 cells were seeded at 1 × 10^5 cells per well in six-well plates and allowed to attach overnight. Cells were collected by trypsinization, washed with PBS, and fixed in 70% ethanol for 15 min on ice. Fixed cells were collected by centrifugation, washed with PBS, and incubated for 30 min in the dark with a staining solution containing 50 μg/mL propidium iodide (Thermo Fisher Scientific, P1304MP) and 50 μg/mL RNase A.

For each sample, 10,000 events were acquired using a BD LSRFortessa flow cytometer operated with FACSDiva software. Debris and doublets were excluded based on light-scatter parameters and PI area versus width. Cell-cycle distributions were determined from DNA-content profiles using the Watson pragmatic model in FlowJo v10.10. Three independent biological experiments were performed for each cell line.

### EdU incorporation and cell-cycle analysis

U2OS WT, ANP32A KO 3.1, and ANP32A KO 3.2 cells were seeded in 60-mm dishes 24 h before labeling. Cells were incubated in pre-warmed culture medium containing 10 μM 5-ethynyl-2′-deoxyuridine (EdU; Cayman Chemical 20518) for 15 min at 37°C. Cells were washed twice with pre-warmed PBS, collected by trypsinization in culture medium, and maintained on ice. After centrifugation at 4°C, cells were washed with PBS and extracted for 5-10 min on ice in CSK buffer containing 25 mM HEPES pH 7.4, 50 mM NaCl, 1 mM EDTA, 3 mM MgCl₂, 300 mM sucrose, 0.5% Triton X-100, and protease inhibitors. Cold PBS containing 1 mg/mL BSA was added, and the cells were collected by centrifugation.

Cell pellets were fixed with 2% paraformaldehyde in PBS for 30 min at room temperature and washed with BD Perm/Wash buffer. Incorporated EdU was detected by incubating cells in 150 μL of freshly prepared click-reaction buffer containing 2 mM CuSO₄, 2 mg/mL sodium L-ascorbate, and 1 μM Alexa Fluor 647 azide (AAT Bioquest) in PBS for at least 30 min at room temperature in the dark. Cells were washed with Perm/Wash buffer and incubated for 20-30 min at 37°C in analysis buffer containing 10 μg/mL propidium iodide, 250 μg/mL RNase A, 0.02% sodium azide, and 1 mg/mL BSA in PBS.

For each sample, 50,000 events within the cell gate were acquired using a BD Accuri flow cytometer. Debris and doublets were excluded, and EdU incorporation and DNA content were analyzed in single cells using FlowJo. Three independent biological experiments were performed for each cell line.

### DNA fiber analysis

Replication-fork progression and stalled-fork protection were assessed using DNA fiber assays. For analysis of unperturbed fork progression, U2OS WT, ANP32A KO 3.1, and ANP32A KO 3.2 cells were seeded at 1.2 × 10^5 cells per well in 12-well plates 24 h before labeling. Cells were sequentially incubated with 100 μM 5-chloro-2′-deoxyuridine (CldU; MilliporeSigma, C6891) for 30 min and 250 μM 5-iodo-2′-deoxyuridine (IdU; MilliporeSigma, I7125) for 30 min, with two washes in pre-warmed medium between the labeling periods. Cells were harvested immediately after the second labeling period.

For stalled fork protection experiments, U2OS WT cells were transfected by reverse transfection using Lipofectamine RNAiMax (Thermo Fisher; 13778150) with siRNAs siCtrl [ sens: GGGUAUCGACGAUUACAAAtt, antisens:UUUGUAAUCGUCGAUACCCtt, Invitrogen], siBRCA1 [ sens: GAAGGAGCUUUCAUCAUUCtt, antisens: GAAUGAUGAAAGCUCCUUCtt) Eurofins Genomics ], or siBRCA2 [ sens: CAGTTGAAATTAAACGGAAtt. Ambion Silencer Select siRNA ], 48 h before the assay. U2OS WT, ANP32A KO 3.1, and ANP32A KO 3.2 cells were analyzed in parallel. Following sequential CldU and IdU labeling as described above, cells were washed twice with pre-warmed medium and treated with 4 mM hydroxyurea (HU; MilliporeSigma, 40046-5GM) for 4 h before collection.

Cells were harvested by trypsinization and resuspended in PBS at 1 × 10^6 cells/mL. Four µl of cell suspension were deposited onto a glass microscope slide and allowed to partially air-dry for 3 min. Twelve µl of lysis buffer containing 200 mM Tris-HCl, pH 7.5, 50 mM EDTA, and 0.5% SDS were added and gently mixed with the cell suspension. After incubation for 5 min, the slides were tilted at an angle of approximately 15° to allow the DNA fibers to spread by gravity. Slides were air-dried for 15 min and fixed for 10 min in a cold solution containing 75% methanol and 25% acetic acid.

Slides were rehydrated in distilled water for 5 min, and DNA was denatured with 2.5 M HCl for 90 min at room temperature. After four washes with PBS, slides were blocked with 5% BSA in PBS for 30 min at 37°C in a humidified chamber. CldU and IdU were detected by overnight incubation at 4°C with rat anti-BrdU antibody (1:250; Abcam, ab6326) and mouse anti-BrdU antibody (1:10; BD Biosciences, 347580), respectively. Slides were washed four times with 0.05% Tween-20 in PBS and once with PBS before incubation for 3 h at 4°C with Alexa Fluor 488-conjugated goat anti-rat IgG (1:100; Thermo Fisher Scientific, A-11006) and Cy3-conjugated sheep anti-mouse IgG F(ab′)₂ fragments (1:100; MilliporeSigma, C2181). Thus, CldU and IdU tracts were visualized in green and red, respectively.

Slides were washed as described above, rinsed sequentially with distilled water and 95% ethanol, air-dried, and mounted using ProLong Diamond Antifade Mountant (Thermo Fisher, P36965). Images were acquired using a ZEISS Axio Observer Z1 microscope equipped with a 63×/1.4-NA oil-immersion Plan-Apochromat objective and an Axiocam 506 monochrome camera. Continuous DNA fibers containing both CldU and IdU tracts were selected, and tract lengths were measured manually using ImageJ. At least 150 fibers were analyzed per biological replicate and condition. Replication-fork velocity was calculated from the conversion of tract length (µm) in kb (conversion factor is 2.59 kb/μm) and labeling time (kb/min).

### Real-time cell proliferation and drug sensitivity assays

Real-time proliferation was measured using the xCELLigence RTCA DP impedance system. Before cell seeding, background impedance was measured in E-Plate VIEW 16 plates containing pre-warmed culture medium (Agilent Technologies, 300601140). U2OS WT, ANP32A KO 3.1, and ANP32A KO 3.2 cells were seeded at 7,500 cells per well and allowed to attach for 24 h.

Cells were then left untreated or exposed continuously to 250 μM hydroxyurea (EMD Millipore; 40046-5GM), 5 μM cisplatin (Cayman Chemical, 13119), 5 μM etoposide (Cayman Chemical, 12092), or 1 μM aphidicolin (Cayman Chemical, 14007). Cellular impedance was measured every 15 min for a total monitoring period of 120 h. Data are presented as mean ± SD from three independent biological experiments.

### Immunoblotting

Cells were washed with PBS and lysed in 1× Laemmli sample buffer. Lysates were sonicated twice for 10 seconds on ice at 25% amplitude using a model FB120110 sonicator (Thermo Fisher Scientific). Protein concentrations were determined using the Pierce BCA Protein Assay Kit (Thermo Fisher Scientific, 23223 and 23224). Depending on the experiment, 20-40 μg of protein per sample were resolved by SDS-PAGE and transferred to nitrocellulose membranes (Bio-Rad, 1620115) using Bjerrum Schafer-Nielsen transfer buffer containing 20% ethanol.

Membranes were incubated with the primary antibodies and corresponding HRP-conjugated secondary antibodies listed in the key resources table. Immunoreactive proteins were detected using Clarity Western ECL Substrate (Bio-Rad, 1705061) and imaged using a ChemiDoc XRS system with Image Lab software (Bio-Rad). Where indicated, band intensities were quantified using Image Lab, and target protein abundance was normalized to GAPDH. Statistical analyses were performed as described under Quantification and statistical analysis.

### Proximity ligation assay

Protein proximity was assessed using the Duolink In Situ Red Starter Kit Mouse/Rabbit (MilliporeSigma, DUO92101). Cells were seeded onto glass coverslips in 12-well plates and subjected to the indicated doxycycline and hydroxyurea (HU) treatments. HU was used at 1 mM for the durations specified for each experiment. Cells were fixed with 4% Paraformaldehyde Electro-Microscopy Grade (Ted Pella, 18505) for 20 min at 4°C and washed twice with PBS. The perimeter of each coverslip was delineated using a Liquid Blocker Super PAP Pen (Electron Microscopy Sciences, 71310). Cells were permeabilized with 0.25% Triton X-100 in PBS for 5 min at 4°C and washed twice with PBS.

Samples were incubated with the kit blocking solution for 1 h at 37°C and subsequently incubated with the appropriate mouse and rabbit primary antibody pairs listed in the key resources table for 2 h at room temperature in the dark. Duolink PLUS and MINUS proximity probes, ligation, and rolling-circle amplification were applied according to the manufacturer’s instructions. Nuclei were stained with DAPI (Biotium, 40011) at 0.5 ng/μL for 5 min, and coverslips were mounted using Immu-Mount (Thermo Fisher Scientific, 9990402).

Images were acquired using a ZEISS LSM 880 laser-scanning confocal microscope operated with ZEN Black software and a 40× oil-immersion objective (2× digital zoom). Image acquisition was performed at a 2048 × 2048 pixel resolution using 12-bit image depth, bidirectional scanning, a pixel dwell time of 0.26 µs, line averaging of 2, and a scan time of 5.03 seconds per optical section. Z-stack images were collected over a 10 µm depth using 21 optical sections separated by 0.5 µm, resulting in a total acquisition time of approximately 1 minute and 52 seconds per field. Maximum-intensity projections (MIP) were generated and analyzed using ZEN Blue version 3.1 and CellProfiler version 4.2.8. Nuclei were segmented using the DAPI signal, and PLA puncta were quantified both throughout the nucleus and within the perinuclear compartment. The perinuclear compartment was defined as a 10-pixel-wide region immediately adjacent to the inner nuclear envelope. This region was generated in CellProfiler by shrinking each segmented nucleus by 10 pixels (*ExpandOrShrinkObjects*) and subtracting the shrunken nucleus from the original nuclear mask (*MaskObjects*). Unless otherwise indicated, 12 fields were analyzed per condition in each experiment. The numbers of analyzed cells and independent biological replicates are reported in the corresponding figure legends.

### DNA Repair Reporter Assay

Homologous recombination (HR) and end-joining (EJ) activity were quantified using the DR-GFP (Gunn and Stark 2012) and EJ5-GFP (Pierce et al. 2001) chromosomal reporter systems, respectively. Both reporters contain I-SceI recognition sites and permit restoration of functional GFP expression following repair of an I-SceI-induced DNA double-strand break through the corresponding repair pathway.

U2OS WT, ANP32A KO 3.1, and ANP32A KO 3.2 cells were transfected with either the DR-GFP or EJ5-GFP reporter construct using Jet-prime (3 μg of DNA for 6 cm petri dish plated at 70% confluence, Polyplus-transfection SA, 10839). Stable integrants were selected with 1 μg/mL puromycin, and individual puromycin-resistant clones were isolated by limiting dilution and expanded. Reporter functionality was evaluated by transient transfection with the pCAG-I-SceI expression plasmid (Addgene, Plasmid #44024). Clones exhibiting a 5 to 10-fold increase in the percentage of GFP-positive cells following I-SceI expression relative to the corresponding no-I-SceI control were retained for subsequent experiments.

Two days after I-SceI transfection, cells were collected and analyzed using a BD LSR Fortessa flow cytometer operated with FACSDiva software. Sequential gates based on forward and side scatter were used to exclude debris and select single cells. GFP-positive gates were established using matched cells that had not received the I-SceI expression plasmid. Reporter activity was expressed as the percentage of GFP-positive singlets. Six independent biological experiments were performed for each genotype and reporter system.

### Laser-Microirradiation

Localized DNA damage was induced by laser microirradiation as previously described (Gaudreau-Lapierre et al. 2018), with the modifications indicated below. U2OS WT cells were seeded at 2.5 × 10⁴ cells per well in µ-Plate 96 Well imaging plates (Ibidi, ibiTreat: #1.5 polymer, cat no 89626) and cultured overnight. Before irradiation, cells were incubated with Hoechst 33342 (Thermo Fisher, H3570) at 10 μg/mL for 15 min and washed twice with phenol red-free DMEM (Wisent, 319-013-CL). The medium was then replaced with fresh phenol red-free DMEM for irradiation.

Defined nuclear regions were irradiated using the 405-nm laser of an Olympus FLUOVIEW FV3000 laser-scanning confocal microscope. Microirradiation was performed using the 405-nm laser at 50% output power in discontinuous mode. For each field of view, 10-12 individual regions of interest (ROIs) were defined and irradiated with a single laser iteration over a 7-pixel line. Images were acquired at 40× magnification using a 1× zoom, 1,024 × 1,024-pixel resolution, and an 8 μs/pixel dwell time. Three independent fields of view were acquired for each experimental condition. Cells were allowed to recover for 5 min, 15 min, 30 min, 90 min, or 180 min and were then fixed with 4% paraformaldehyde in PBS. ANP32A and γH2AX were subsequently detected by immunofluorescence as described below.

The proportions of cells displaying ANP32A-γH2AX colocalization as well as colocalization events located at the nuclear periphery were determined manually. The numbers of analyzed cells and independent experiments are reported in the corresponding figure legend.

### Immunofluorescence staining and confocal microscopy

U2OSFT, U2OS WT, ANP32A KO 3.1, and ANP32A KO 3.2 cells were grown on glass coverslips to approximately 60% confluence. Laser-microirradiated cells were processed directly in 96-well imaging plates. Cells were fixed with 4% paraformaldehyde for 20 min at 4°C, washed with PBS, and permeabilized with 0.15% Triton X-100 in PBS for 5 min at 4°C. After washing, samples were blocked with 3% BSA in PBS for 20 min at 4°C.

Samples were incubated for 2 h at room temperature with the primary antibodies listed in the key resources table. After washing, cells were incubated for 1 h at room temperature with Alexa Fluor 488-conjugated goat anti-mouse IgG (1:800; Thermo Fisher Scientific, A-11001) or Alexa Fluor 546-conjugated goat anti-rabbit IgG (1:800; Thermo Fisher Scientific, A-11010), as appropriate. Nuclei were stained with DAPI (Biotium, 40011) at 1 μg/mL for 5 min. Coverslips were mounted using Immu-Mount (Thermo Fisher Scientific, 9990402).

Images were acquired using a ZEISS LSM 880 laser-scanning confocal microscope controlled by ZEN Black software with a 40× oil-immersion objective, a 2× digital zoom, and a resolution of 1,024 × 1,024 pixels. Identical acquisition settings were used for all conditions within an experiment. Images were processed and quantified using ZEN Blue version 3.1 and Fiji Image J version 2.16.0 . The numbers of fields, cells, and independent biological experiments analyzed are reported in the corresponding figure legends.

### AirID proximity labeling

#### Twenty-four-hour ANP32A proximity labeling

U2OSFT cells stably expressing AirID-3×Myc-ANP32A or the AirID-3×Myc negative control were cultured in 100-mm dishes to approximately 40%-50% confluence. Expression of the AirID constructs was induced with 2.5 μg/mL doxycycline (Takara Bio (formerly Clontech Laboratories), 631311) for 48 h. During the final 24 h of induction, cells were supplemented with 50 μM biotin (Iris Biotech, LS-3500.0250) and either left untreated or exposed to 1 mM hydroxyurea (HU) (when applicable; EMD Millipore; 40046-5GM). Three independent biological replicates were prepared for each cell line and treatment condition. Cells were harvested by trypsinization and collected by centrifugation at 1,500 × g for 5 min at 4°C. Cell pellets were subsequently processed for streptavidin affinity purification as described below.

#### Time-resolved ANP32A proximity labeling

For time-resolved analysis, U2OSFT cells stably expressing AirID-3×Myc-ANP32A were cultured and induced with 2.5 μg/mL doxycycline for 48 h as described above. Cells were left untreated or exposed to 1 mM HU for 3 h, 6 h, 12 h, or 24 h. Biotin was added to a final concentration of 50 μM during only the final 3 h preceding collection. Thus, for the 3 h HU condition, HU and biotin were added simultaneously, whereas biotin was added during the final 3 h of the longer HU treatments. Untreated cells were similarly labeled with biotin for 3 h and served as the 0 h reference condition. One 100-mm dish was prepared per condition for each biological replicate, and three independent biological replicates were analyzed. Cells were harvested by trypsinization and centrifuged at 1,500 × g for 5 min at 4°C.

### Sample preparation for mass spectrometry

#### Streptavidin affinity purification of biotinylated proteins

Cell pellets were resuspended in freshly prepared lysis buffer containing 8 M urea (Sigma, U5128-5KG), 50 mM HEPES (Wisent, 330-050-EL) pH 7.4, 1 mM phenylmethylsulfonyl fluoride (PMSF) (Themo Fisher Scientific, 71105GM), and 1 mM dithiothreitol (DTT) (Life Technologies, R0862). Lysates were transferred to low protein-binding tubes and sonicated on ice for two 1-min cycles using 10 s on/10 s off pulses at 25% amplitude with a model FB120110 sonicator. Insoluble material was removed by centrifugation at 16,500 × g for 10 min at 4°C.

Protein concentrations were determined using the Pierce BCA Protein Assay Kit (Themo Fisher Scientific, PI23223, PI23224). Aliquots were retained for immunoblot verification of AirID-fusion-protein expression and proximity-dependent biotinylation. Each sample was normalized to 0.5 mg of total protein in 1 mL of lysis buffer and incubated with 25 μL of prewashed high-performance streptavidin Sepharose beads (MilliporeSigma, GE17-5113-01). Samples were incubated for 2 h at room temperature with end-over-end rotation.

Beads were collected by centrifugation at 1,000 × g for 5 min at room temperature, and the supernatants were carefully removed. The beads were washed five times with 8 M urea and 50 mM HEPES, pH 7.4. Following the final wash, beads were transferred to new low-protein-binding tubes (Sarstedt, 72.701.600) and processed for on-bead tryptic digestion.

#### On-bead tryptic digestion and peptide desalting

All solutions were prepared using MS-grade water (Fluka Analytical, 39253). Streptavidin Sepharose beads were washed five times with 20 mM ammonium bicarbonate (MilliporeSigma, 09830), followed by an additional wash with 20 mM ammonium bicarbonate containing 1 mM biotin to saturate residual streptavidin-binding sites.

For on-bead digestion, proteins were reduced in 50 μL of 20 mM ammonium bicarbonate containing 10 mM DTT for 30 min at 60°C with agitation at 1,250 rpm. Samples were then alkylated by adding 50 μL of 20 mM ammonium bicarbonate containing 15 mM chloroacetamide (MilliporeSigma, C0267), followed by incubation for 1 h at room temperature in the dark with continuous agitation. Excess chloroacetamide was quenched by adding DTT to a final concentration of 15 mM.

Proteins were digested overnight at 37°C with 1 μg of MS-grade trypsin (Thermo Fisher Scientific, 90058), corresponding to a final trypsin concentration of 10 μg/mL. Following digestion, samples were acidified with formic acid (Thermo Fisher Scientific, A11750) to a final concentration of 1%. Beads were collected by centrifugation at 2,000 × g for 3 min, and the peptide-containing supernatants were transferred to fresh low protein-binding tubes.

The beads were subsequently resuspended in 100 μL of 60% acetonitrile (VWR, CAAX0156-1) and 0.1% formic acid and incubated for 5 min at room temperature with agitation at 1,250 rpm. The resulting eluates were collected and pooled with the corresponding initial peptide fractions. Pooled samples were dried to completion in a centrifugal evaporator maintained at 60°C and reconstituted in 300 μL of 0.1% trifluoroacetic acid (Thermo Fisher Scientific, A11650).

Peptides were desalted using C18 ZipTip pipette tips (VWR, 88777). Before sample loading, the tips were conditioned three times with acetonitrile and equilibrated three times with 0.1% trifluoroacetic acid. Each 100 μL sample aliquot was loaded using 10 repeated aspiration-dispensing cycles. Bound peptides were washed three times with 0.1% trifluoroacetic acid and eluted into fresh low protein-binding tubes using 50% acetonitrile and 1% formic acid. The loading and elution procedures were repeated three times to process the complete sample volume.

Eluted peptides were dried in a centrifugal evaporator at 60°C and reconstituted in 30 μL of 1% formic acid. Peptide concentrations were estimated using a NanoDrop 2000c spectrophotometer (Thermo Fisher Scientific, ND-2000C). Samples were transferred to glass autosampler vials (Thermo Fisher Scientific, 6PRV11-03FIVP) and stored at −20°C until LC-MS/MS analysis.

### LC-MS/MS analysis

#### 24-h ANP32A proxeome

Peptides from the 24 h ANP32A proxeome experiment were analyzed using a nanoElute 2 UHPLC system coupled to a timsTOF Pro ion-mobility mass spectrometer equipped with a CaptiveSpray nanoelectrospray-ionization source (Bruker Daltonics). For each sample, 500 ng of peptides were loaded onto an Acclaim PepMap 100 C18 trap column (0.3 × 5 mm; Thermo Fisher Scientific) and separated on a PepSep analytical column packed with 1.9 μm C18 particles (75 μm × 25 cm; Bruker Daltonics).

Peptides were separated over a 120 min gradient from 5% to 37% acetonitrile in 0.1% formic acid at a flow rate of 400 nL/min. Data were acquired in diaPASEF mode using a target intensity of 20,000 and an intensity threshold of 2,500. The acquisition method used a 100-ms TIMS ramp, 27 mass steps with a width of 50 Da spanning m/z 114-1414, a duty cycle of 1.27 s, and a collision energy of 42 eV. The acquisition scheme was designed to follow the diagonal distribution of doubly and triply charged peptide precursors in m/z-ion-mobility space.

#### Time-resolved ANP32A proxeome

Peptides from the time-resolved ANP32A proxeome experiment were analyzed using an Orbitrap Astral mass spectrometer (Thermo Fisher Scientific; RRID: SCR_026205) coupled to a Vanquish Neo UHPLC system and operated with a nanoelectrospray interface in positive-ion mode. For each sample, 100 ng was injected.

Mobile phase A consisted of 0.1% formic acid in water, and mobile phase B consisted of 0.1% formic acid in 80% acetonitrile. Peptides were first loaded onto a PepMap Neo C18 trap column (5 μm, 300 μm × 5 mm) and subsequently separated on a fused-silica analytical column (100 μm internal diameter × 150 mm) packed in-house with 1.9 μm ReproSil-Pur C18 resin with a pore size of 100 Å (Dr. Maisch GmbH).

Peptides were separated at 500 nL/min using a linear gradient from 5% to 35% mobile phase B over 18.5 min. The spray voltage was set to 2.5 kV, and the ion-transfer-tube temperature was maintained at 300°C. MS1 and DIA spectra were acquired over an m/z range of 380-980. DIA acquisition used 2 Da isolation windows and higher-energy collisional dissociation at a normalized collision energy of 25%. Full-scan Orbitrap spectra were acquired at a resolution of 240,000, and real-time internal mass calibration was performed using a lock-mass strategy. Raw data were acquired using Xcalibur version 4.7.

### Proteomics data processing and statistical analysis

#### Protein identification and quantification using DIA-NN

Raw files from the 24 h ANP32A proxeome experiment were analyzed using DIA-NN version 2.2.0 against the UniProt human proteome database downloaded on March 10, 2024 (83,132 entries). Raw files from the time-resolved ANP32A proxeome experiment were analyzed using DIA-NN version 2.3.0 against the UniProt human proteome database downloaded on May 18, 2025 (83,180 entries).

Both datasets were processed using a library-free workflow. In silico FASTA digestion and deep-learning-based prediction of MS/MS spectra, retention times, and ion-mobility values were enabled. Trypsin was specified as the proteolytic enzyme, allowing up to one missed cleavage. Peptides of 7-30 amino acids with precursor charge states of +2 to +4 were considered. Carbamidomethylation of cysteine was specified as a fixed modification, and N-terminal methionine excision was enabled. Precursor- and fragment-ion mass tolerances were set to 20 ppm, and the precursor-level false-discovery rate was controlled at 1%. Match Between Runs, Unrelated Runs, and MaxLFQ were enabled for identification transfer and label-free quantification.

#### Protein-level processing and differential-abundance analysis

Three independent biological replicates were analyzed for each condition. Protein-level quantitative data were imported into ProStaR version 1.38.1 (Wieczorek et al. 2017). Proteins identified with fewer than two unique peptides were excluded. Missing values were classified as partially observed values (POVs), corresponding to proteins quantified in only a subset of replicates, or missing in an entire condition (MEC). Proteins identified with 3 POV of the 3 biological replicas in at least one condition were kept while the ones founds with ≥2 POV in at least one condition were deleted. MEC were then removed for both conditions. Remaining missing values were imputed using the DetQuantile method with a 1% quantile and a factor of 0.2. Variance-stabilizing normalization was subsequently applied. Proteins were retained for statistical analysis when quantified in at least two of the three biological replicates. Differential abundance was assessed using a Limma moderated t test. P values were adjusted for multiple testing using the Benjamini-Hochberg procedure with π₀ = 1. Proteins with a raw p value ≤ 1 × 10⁻³ and an adjusted p value ≤ 0.05 were considered differentially abundant.

#### SAINTexpress analysis of the 24 h ANP32A proxeome

Data from the 24 h ANP32A proxeome experiment were additionally analyzed using SAINTexpress version 3.6.1. AirID-3×Myc-ANP32A was treated as the bait, and the AirID-3×Myc samples were used to estimate nonspecific background labeling. SAINTexpress (Choi et al. 2011; Teo et al. 2014) used protein-level MS intensities from the three biological replicates to calculate the probability of each bait-prey association. High-confidence proximal proteins were defined using a SAINT score ≥ 0.8 and a Bayesian false-discovery rate (BFDR) <5%. Fold enrichment relative to the AirID-3×Myc control was also calculated, and most retained proteins displayed at least twofold enrichment. Complete results are provided in Table S2. Functional enrichment of the high-confidence proximal proteins identified under basal and HU-treated conditions was performed using STRING version 12.0 (Szklarczyk et al. 2023) based on Gene Ontology (GO) Molecular Function and subcellular localization annotations. Enrichment visualization was generated using the STRING enrichment analysis tool with a false discovery rate (FDR) threshold of ≤ 0.05. GO terms were displayed without term merging, using a minimum signal of 0.01, a minimum enrichment strength of 0.01, and a minimum network count of two proteins per term. Functional terms were ranked according to the enrichment signal and grouped using a similarity threshold of ≥ 0.8.

#### Analysis of the time-resolved ANP32A proxeome

The time-resolved ANP32A proxeome was processed in ProStaR using the filtering, imputation, normalization, and statistical procedures described above. Each HU-treatment duration was compared with the untreated 0 h reference using three independent biological replicates per condition. Proteins with a log₂ fold change ≤ −1 or ≥ 1 that also passed the ProStaR statistical-significance threshold with a FDR threshold of ≤ 5% were considered significantly depleted or enriched, respectively, in the ANP32A proxeome. Complete results are provided in Table S3. Significant protein sets across the HU treatment time points were compared using InteractiVenn (Heberle et al. 2015), and functional enrichment analyses were performed using ShinyGO (Ge et al. 2020) based on GO Biological process databases with a FDR threshold of ≤ 0.05.

#### Quantification and statistical analysis

Unless otherwise indicated, statistical analyses were performed using GraphPad Prism v11.0.2. Data are presented as mean ± SD, and the number of independent biological replicates, analyzed cells, images, or DNA fibers is specified in the corresponding figure legends. A two-sided p value < 0.05 was considered statistically significant. Exact p values are reported whenever available; ns denotes a non-significant difference. Proteomics-specific filtering, imputation, normalization, differential abundance testing, and multiple testing correction are described in the preceding section.

Three independent experiments of crystal-violet proliferation measurements, cell-cycle distributions determined by propidium-iodide staining, EdU-incorporation measurements, and xCELLigence proliferation curves were analyzed using two-way ANOVA followed by Dunnett’s multiple-comparisons test. Comparisons were performed between each ANP32A knockout clone and the corresponding U2OS WT condition.

Immunoblot band intensities were quantified using Image Lab and normalized to GAPDH. Cyclin D1 abundance was quantified from six independent biological experiments, whereas RB and E2F1 abundance was quantified from three independent biological experiments. Comparisons between each ANP32A-knockout clone and U2OS WT cells were performed using Welch’s t-test between group U2OS WT VS Group U2OS KO 3.1 and 3.2.

For PLA experiments comparing untreated and HU-treated cells at a single time point, statistical significance was assessed using a two-tailed Mann-Whitney U test. PLA experiments containing more than two conditions, including the HU time courses, were analyzed using a Kruskal-Wallis test. PLA puncta were quantified throughout the nucleus and within the perinuclear compartment. The numbers of analyzed cells, fields, and biological replicates are provided in the corresponding figure legends.

Replication-fork velocities were compared between U2OS WT and ANP32A-knockout cells using two-way ANOVA followed by Šidák’s multiple-comparisons test. Stalled fork-protection measurements, expressed as IdU/CldU tract-length ratios, were analyzed two-way ANOVA followed by Šidák’s multiple-comparisons test. At least 150 DNA fibers were measured per biological replicate and condition.

DR-GFP homologous-recombination and EJ5-GFP end-joining reporter activities were compared among U2OS WT, ANP32A KO 3.1, and ANP32A KO 3.2 cells using a Kruskal-Wallis test followed by multiple comparisons using U2OS WT cell line as control. Six independent biological experiments were performed for each reporter system.

The laser microirradiation experiment shown in Figure 6C and 6D was performed once, with at least 89 cells analyzed per time point, and is therefore presented descriptively without inferential statistical testing. ANP32A-γH2AX PLA measurements from three independent biological experiments were analyzed using a Kruskal-Wallis test followed by multiple comparisons using untreated condition as control.

## RESULTS

### AirID mapping identifies a replication stress-regulated ANP32A proximal proteome

To define the molecular environment of ANP32A and determine how it changes during replication stress, we generated isogenic U2OS Flp-In T-REx cells expressing AirID-3×Myc-ANP32A or AirID-3×Myc as a background labeling control (Kido et al. 2020). Expression was induced with doxycycline for 48h, and cells were supplemented with biotin during the final 24 h, with or without 1 mM hydroxyurea (HU). Biotinylated proteins were then affinity purified and analyzed by data-independent acquisition mass spectrometry (DIA-MS) (Figure 1A). Immunoblotting confirmed expression of the AirID constructs and robust proximity biotinylation under both untreated and HU-treated conditions (Figure 1B and Figures S1A and S1B). Immunofluorescence microscopy further showed that AirID-3×Myc-ANP32A localized predominantly to the nucleus, consistent with the localization of endogenous ANP32A (Figure 1C). These results established an inducible system for comparing the ANP32A proximal proteome under basal and replication-stress conditions.

**Figure 1:**
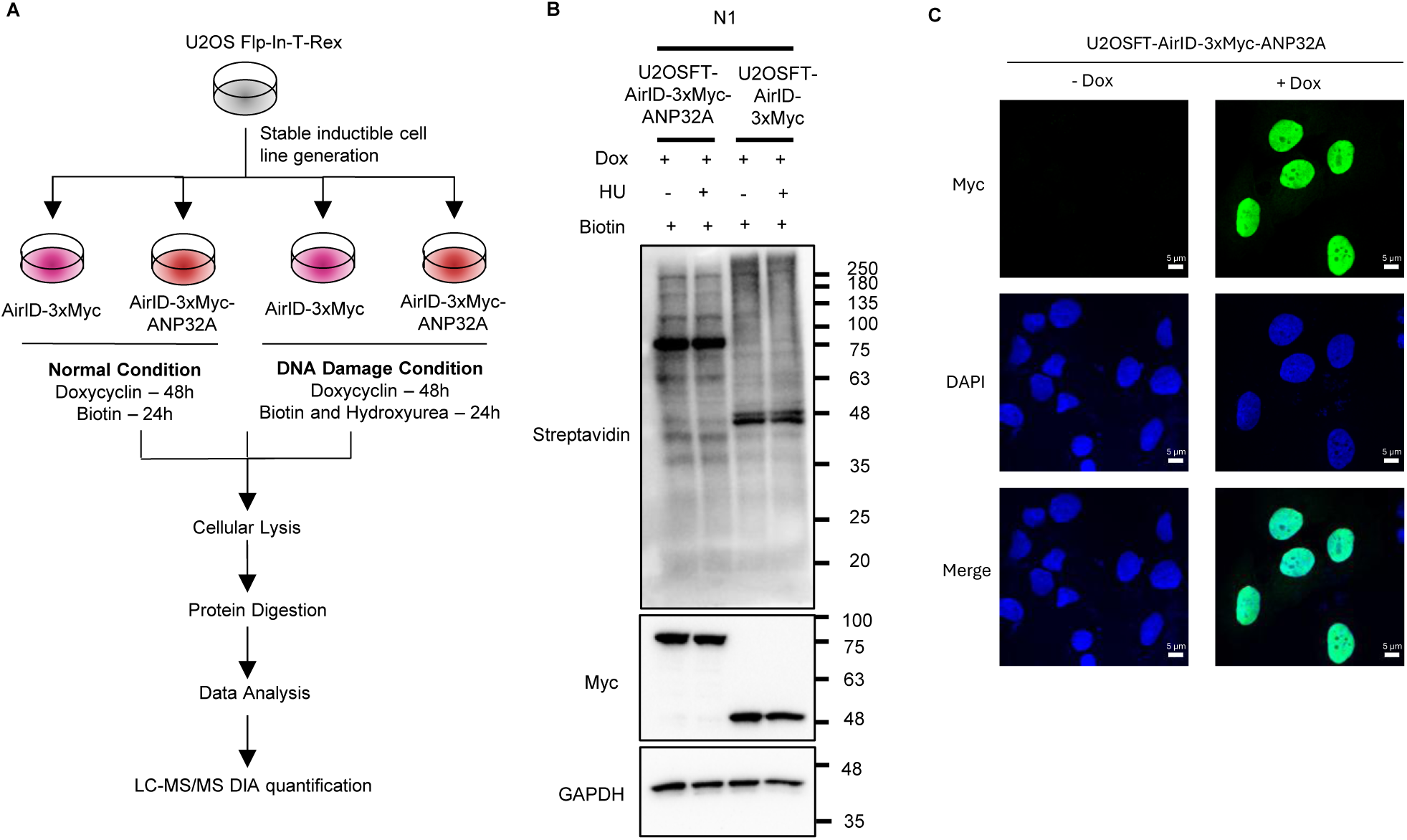
Experimental design and validation of the inducible AirID proximity-labeling system. (**A**) Experimental workflow for defining the ANP32A proximal proteome under basal and replication stress conditions. U2OS Flp-In T-REx (U2OSFT) cells stably expressing AirID-3×Myc-ANP32A or AirID-3×Myc were induced with doxycycline for 48 h. Biotin was added during the final 24 h, with or without 1 mM hydroxyurea (HU), followed by cell lysis, streptavidin affinity purification, protein digestion, and DIA-MS analysis. (**B**) Immunoblot validation of AirID-3×Myc-ANP32A and AirID-3×Myc expression and proximity biotinylation under untreated and HU-treated conditions. Biotinylated proteins were detected using streptavidin-HRP, AirID constructs using an anti-Myc antibody, and GAPDH as a loading control. Data are from biological replicate N1; additional replicates are shown in Figure S1. (**C**) Immunofluorescence analysis of AirID-3×Myc-ANP32A expression and localization before and after doxycycline induction. Myc is shown in green and DNA was counterstained with DAPI (blue). Scale bars, 5 μm.

High-confidence ANP32A-proximal proteins were identified using SAINTexpress relative to the AirID-only control. Under basal conditions, four proteins met the significance criteria: ANP32A itself, SGF29, ASF1B, and VPS35L (Figure 2A and Table S2). This restricted proximal proteome was enriched for histone-binding and chromatin-associated functions, consistent with established roles of ANP32A in chromatin regulation (Figure 2B). Following 24 h of HU treatment, the ANP32A proximal proteome expanded to 25 high-confidence proteins, including the NHEJ factors XRCC5/Ku80 and XRCC6/Ku70; the 14-3-3 proteins YWHAE and YWHAZ; and BAG2, WDR36, YEATS2, YARS1, and YARS2 (Figure 2C and Table S2). Functional enrichment analysis identified processes related to chromatin regulation, RNA metabolism, protein quality control, and DNA repair, including NHEJ (Figure 2D). Thus, prolonged replication stress substantially remodels the molecular environment surrounding ANP32A.

**Figure 2:**
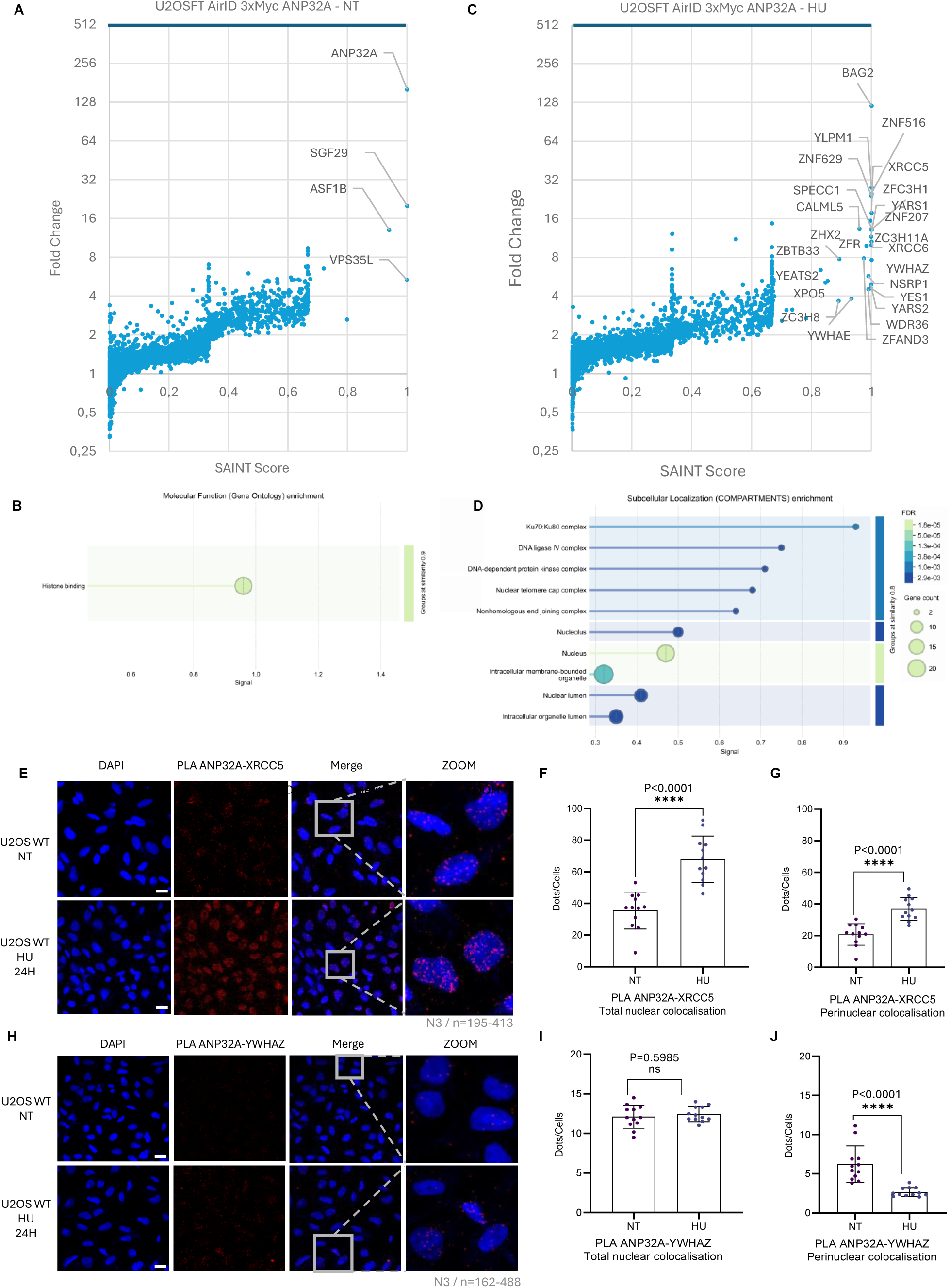
Replication stress remodels the ANP32A proximal proteome. (**A**) SAINTexpress analysis of proteins identified in proximity to AirID-3×Myc-ANP32A relative to the AirID-3×Myc control under untreated conditions. Proteins with a SAINT score ≥0.8 were considered high-confidence proximal proteins. (**B**) Gene Ontology molecular function enrichment analysis of the high-confidence proteins identified under untreated conditions. Circle size represents gene count, and color represents the false discovery rate (FDR). (**C**) SAINTexpress analysis of ANP32A-proximal proteins following treatment with 1 mM HU for 24 h. (**D**) Gene Ontology cellular component enrichment analysis of the high-confidence HU-associated proximal proteins. Circle size represents gene count, and color represents FDR. (**E**) Representative proximity ligation assay (PLA) images showing endogenous ANP32A-XRCC5 proximity in untreated U2OS WT cells and following 24 h of HU treatment. PLA puncta are shown in red and nuclei in blue. (**F** and **G**) Quantification of ANP32A-XRCC5 PLA puncta throughout the nucleus (**F**) and at the nuclear periphery (**G**). (**H**) Representative images of endogenous ANP32A-YWHAZ PLA under the same conditions. (**I** and **J**) Quantification of ANP32A-YWHAZ PLA puncta throughout the nucleus **(I)** and at the nuclear periphery (**J**). Data are mean ± SD from three independent biological replicates (N=3), with 12 images analyzed per condition containing 195-413 cells for XRCC5 or 162-488 cells for YWHAZ. Statistical significance was assessed using two-tailed Mann-Whitney tests. Exact P values are indicated; ns, not significant. Scale bars, 20 μm.

To independently validate selected HU-associated proximal proteins, we examined the proximity of endogenous ANP32A to XRCC5 and YWHAZ by proximity ligation assay (PLA) in untreated and HU-treated U2OS cells (Figures 2E and 2H). HU treatment increased ANP32A-XRCC5 proximity from an average of 35.5 to 67.9 puncta per nucleus (Figure 2F). Perinuclear signals similarly increased from 20.7 to 36.8 puncta per cell (Figure 2G). By contrast, total ANP32A-YWHAZ proximity was unchanged following HU treatment, with averages of 12.1 and 12.4 puncta per nucleus in untreated and treated cells, respectively (Figure 2I). However, perinuclear ANP32A-YWHAZ signals decreased from 6.2 to 2.6 puncta per cell (Figure 2J), suggesting spatial redistribution of this association during replication stress. These orthogonal analyses validate the proximity of ANP32A to both candidates while specifically supporting an HU-induced increase in its association with XRCC5.

As an additional specificity control, ANP32A-XRCC5 PLA was performed in U2OS WT cells and two independent ANP32A-knockout clones (KO 3.1 and KO 3.2) described in detail below. Compared with WT cells, PLA signals were markedly reduced in both knockout clones, indicating that the puncta detected in WT cells depend on the presence of ANP32A (Figure S2). This genetic control supports the specificity of the ANP32A-XRCC5 PLA readout and its use as an orthogonal validation of proximity relationships identified by AirID.

### ANP32A proximity networks exhibit distinct temporal dynamics during replication stress

To resolve how the molecular environment surrounding ANP32A changes as replication stress develops, we performed time-resolved AirID proximity labeling in U2OSFT cells expressing AirID-3×Myc-ANP32A. Cells were exposed to 1 mM HU for 0 h, 3 h, 6 h, 12 h, or 24 h, with biotin added during the final 3 h before harvest at each time point (Figure 3A). This design provided matched 3 h labeling windows across all conditions. Immunoblotting confirmed AirID-3×Myc-ANP32A expression and robust proximity biotinylation throughout the time course, while immunofluorescence showed that the fusion protein remained predominantly nuclear during HU exposure (Figure 3B and Figures S3A-S3C).

**Figure 3:**
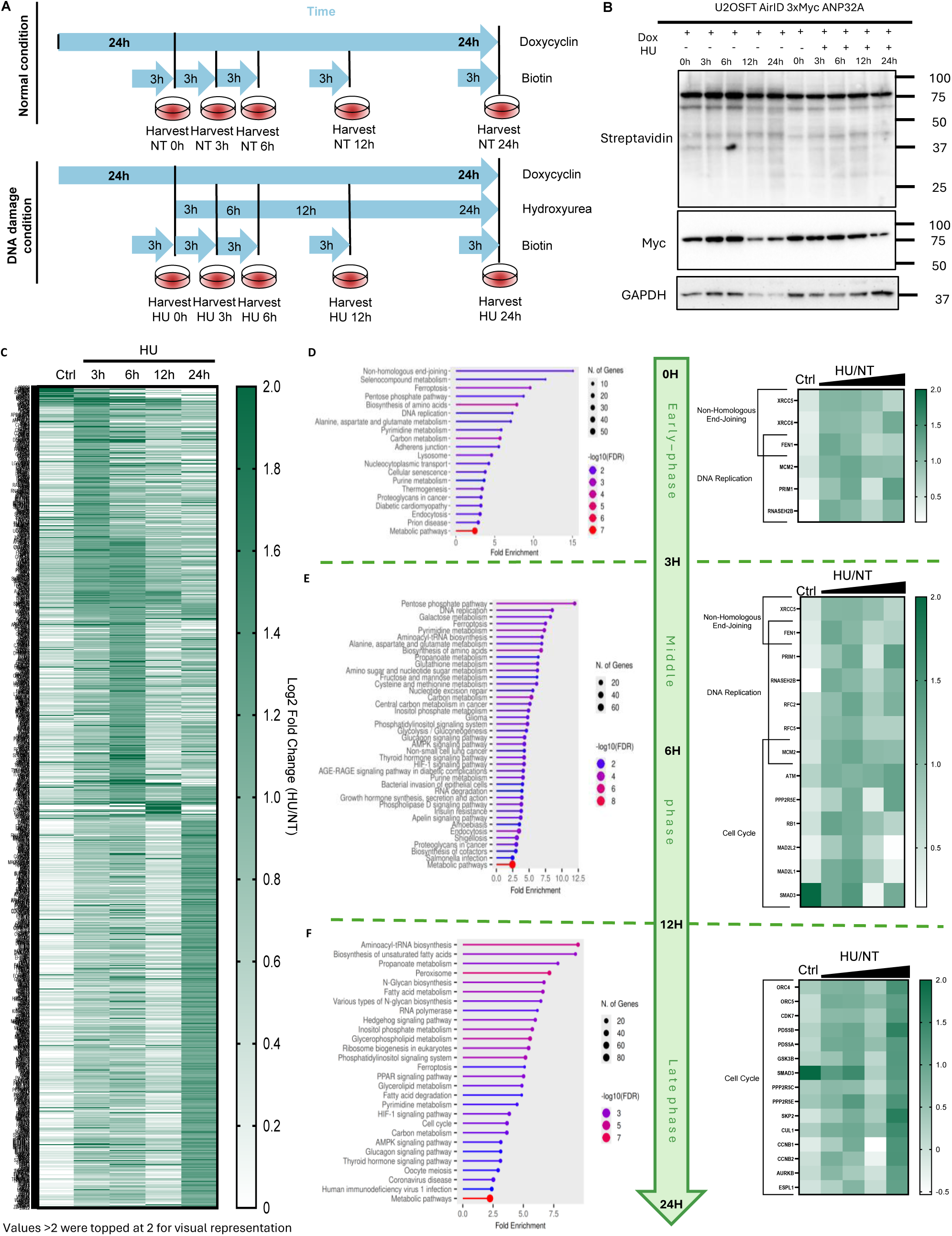
Time-resolved proximity proteomics reveals phased remodeling of the ANP32A proximal proteome. (**A**) Experimental design for time-resolved AirID proximity labeling. U2OSFT cells expressing AirID-3×Myc-ANP32A were induced with doxycycline for 48 h and exposed to 1 mM HU for 0 h, 3 h, 6 h, 12 h, or 24 h. Biotin was added during the final 3 h before each harvest, providing matched labeling windows across all conditions. Biotinylated proteins were purified and analyzed by DIA-MS. (**B**) Immunoblot validation of AirID-3×Myc-ANP32A expression and proximity biotinylation in matched untreated and HU-treated samples collected at the indicated time points. Biotinylated proteins were detected using streptavidin-HRP, the fusion protein using an anti-Myc antibody, and GAPDH as a loading control. (**C**) Heatmap showing the relative abundance of proteins enriched in proximity to ANP32A at one or more time points during HU treatment. Values represent log2 fold changes between HU-treated and matched untreated samples; values greater than 2 were capped at 2 for visualization. (**D-F**) Functional enrichment analyses and heatmaps of selected proteins associated with the early, 0-3 h (**D**); intermediate, 3-12 h (**E**); and late, 12-24 h (**F**), phases of the response. Dot size represents the number of genes assigned to each term, and color represents -log10(FDR).

Biotinylated proteins were isolated and quantified by DIA-MS, and each HU time point was compared with untreated cells to identify stress-dependent changes in ANP32A proximity (Figures S4A-S4E and Table S3). Overlap analysis revealed both shared and time-specific enriched or depleted proteins (Figures S4F and S4G), while visualization of enriched proteins across the complete time course revealed distinct temporal patterns (Figure 3C). During the early response to HU (0-3 h), ANP32A-proximal proteins were enriched for DNA replication and repair functions, including the NHEJ factors XRCC5/Ku80 and XRCC6/Ku70, together with FEN1, MCM2, PRIM1, and RNASEH2B (Figure 3D). At intermediate time points (3-12 h), replication and repair proteins remained prominent, while checkpoint and cell-cycle regulators, including ATM, MAD2L1, PPP2R5E, RB1, and SMAD3, became increasingly represented (Figure 3E). During prolonged HU exposure (12-24 h), the proximal proteome shifted toward cell-cycle and mitotic regulators, including ORC4, ORC5, CDK7, CCNB1, CCNB2, CUL1, AURKB, and ESPL1 (Figure 3F). Thus, the ANP32A proximal proteome undergoes phased remodeling during replication stress, progressing from an early replication and repair-associated network toward checkpoint, cell-cycle, and mitotic regulatory networks.

To validate selected kinetic profiles at the endogenous protein level, we measured ANP32A proximity to XRCC5 and FEN1 by PLA over the same HU time course (Figures 4A and 4D). ANP32A-XRCC5 proximity increased rapidly from an average of 42.6 puncta per nucleus in untreated cells to 136.2, 120.8, and 133.5 puncta after 3 h, 6 h, and 12 h of HU exposure, respectively (Figure 4B). At 24 h, the signal declined toward basal levels, reaching 55.0 puncta per nucleus. Perinuclear signals followed a similar temporal pattern (Figure 4C), supporting a transient increase in ANP32A-XRCC5 proximity during the early and intermediate phases of replication stress.

**Figure 4:**
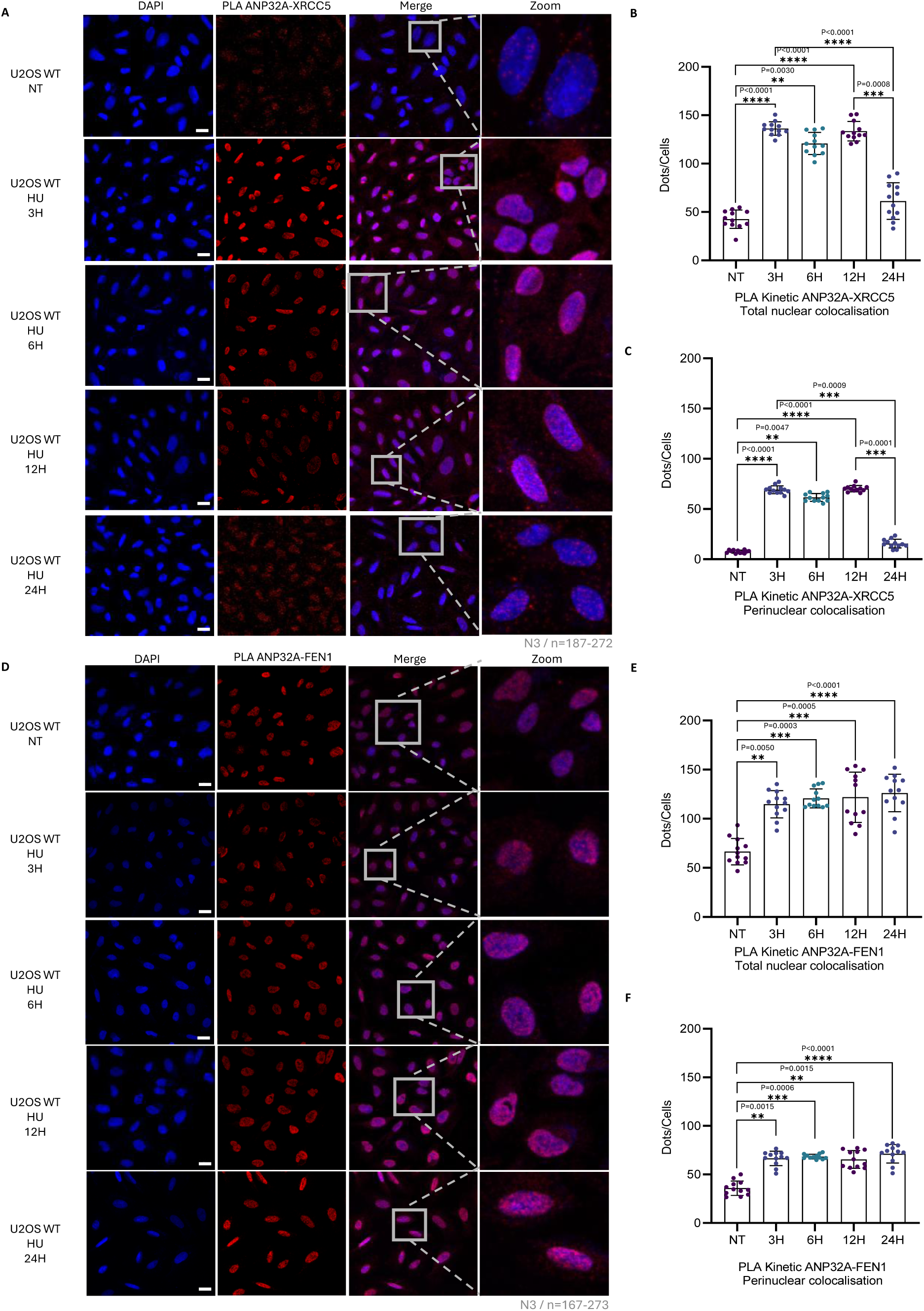
ANP32A-XRCC5 and ANP32A-FEN1 proximity profiles exhibit distinct temporal dynamics during replication stress. (**A** and **D**) Representative PLA images showing endogenous ANP32A proximity to XRCC5 (**A**) or FEN1 (**D**) in untreated U2OS WT cells and after 3 h, 6 h, 12 h, or 24 h of HU treatment. PLA puncta are shown in red and nuclei in blue. **(B** and **E**) Quantification of ANP32A-XRCC5 (**B**) and ANP32A-FEN1 (**E**) PLA puncta throughout the nucleus. (**C** and **F**) Quantification of ANP32A-XRCC5 (**C**) and ANP32A-FEN1 **(F**) PLA puncta at the nuclear periphery. Data are mean ± SD from three independent biological replicates (N=3), with 12 images analyzed per condition containing 187-272 cells for XRCC5 or 167-273 cells for FEN1. Statistical significance was assessed using a Kruskal-Wallis test followed by Dunn’s multiple-comparisons test. Only significant comparisons and their exact P values are indicated. Scale bars, 20 μm.

ANP32A-FEN1 proximity displayed a distinct profile. Relative to 66.5 puncta per nucleus in untreated cells, PLA signals increased to 114.7, 120.7, 121.6, and 126.1 puncta after 3 h, 6 h, 12 h, and 24 h of HU treatment, respectively (Figure 4E). Perinuclear signals showed a comparable sustained increase (Figure 4F). Thus, whereas ANP32A proximity to XRCC5 was transient, its proximity to FEN1 remained elevated throughout the replication stress response. Although FEN1 did not meet the enrichment threshold in the separate 24 h AirID experiment, the PLA results support sustained endogenous proximity between ANP32A and FEN1.

To determine whether PLA also recapitulated proteins that were not enriched in the ANP32A proxeome, we examined ORC1, CHK1, LIG4, and RAD52. None of these proteins met the enrichment criteria in the AirID analysis, and consistent with these proteomic results, PLA detected only low levels of proximity to endogenous ANP32A under both untreated and HU-treated conditions (Figures S5A-S5L). These orthogonal negative results corroborate the lack of enrichment observed by AirID and further support the specificity of the stress-regulated ANP32A proximity network, including its associations with XRCC5 and FEN1. Together, the time-resolved proteomic and PLA analyses reveal selective and temporally distinct ANP32A-associated networks spanning DNA replication, NHEJ, and cell-cycle regulation.

### ANP32A loss impairs proliferation and G1/S progression

Because the time-resolved proxeome linked ANP32A to multiple cell-cycle regulators, we investigated whether ANP32A contributes to proliferation and cell-cycle progression under unperturbed conditions. Using a single sgRNA, we generated two independently isolated ANP32A-knockout U2OS clones, KO 3.1 and KO 3.2 (Figure 5A). Immunoblotting confirmed the loss of ANP32A expression in both clones (Figure 5B). When equal numbers of WT and knockout cells were seeded and monitored by crystal violet staining, both ANP32A-knockout clones exhibited reduced cell accumulation throughout the assay (Figures 5C and 5D). Thus, ANP32A loss compromises the proliferative capacity of U2OS cells.

**Figure 5:**
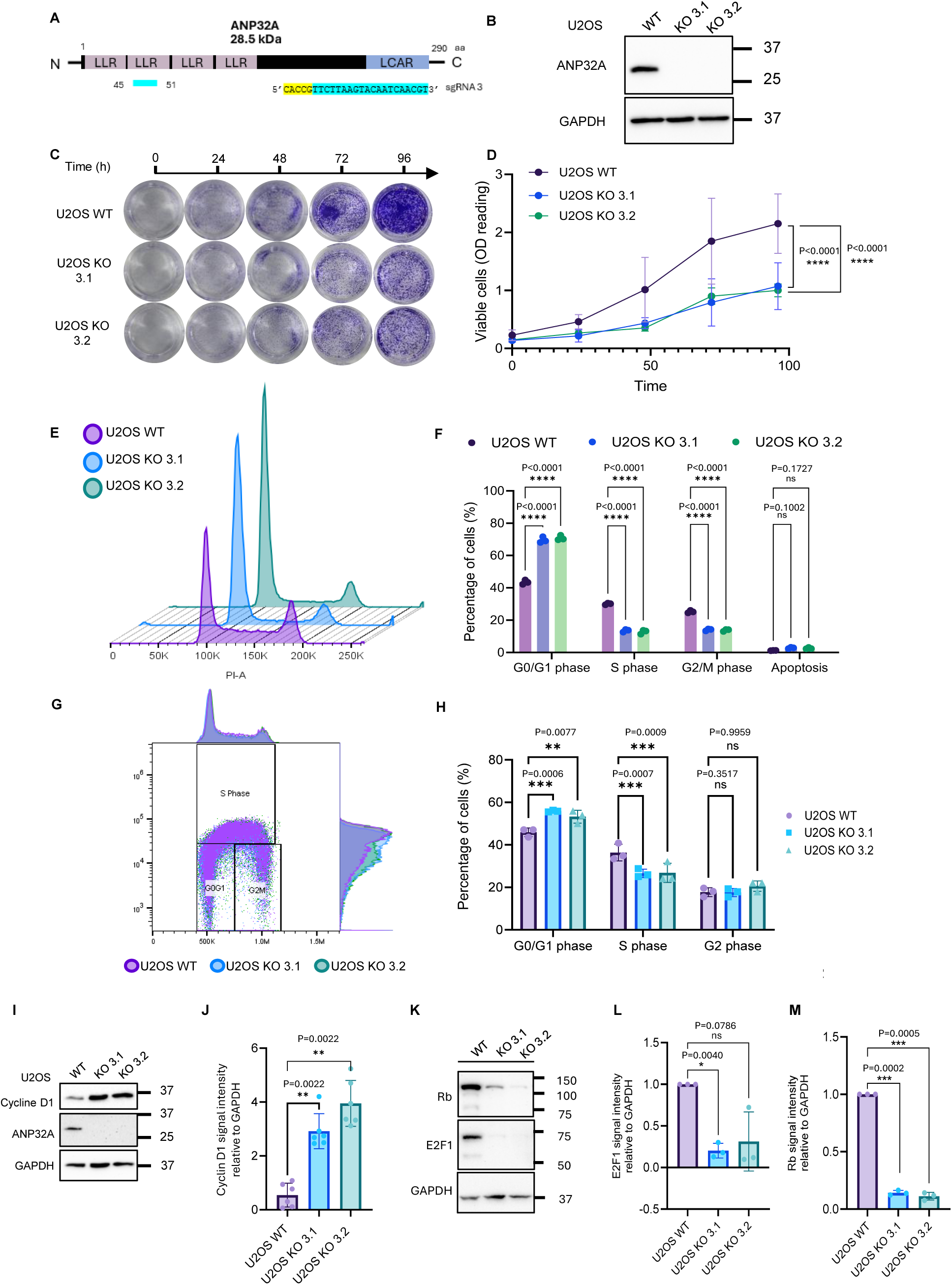
ANP32A loss impairs proliferation and G1/S progression. (**A**) Schematic representation of ANP32A showing its leucine-rich repeats (LRRs), low-complexity acidic region (LCAR), and the position and sequence targeted by sgRNA 3. (**B**) Immunoblot confirming loss of ANP32A expression in independently isolated U2OS ANP32A-knockout clones 3.1 and 3.2. GAPDH was used as a loading control. (**C**) Representative crystal-violet staining of equal numbers of U2OS WT, ANP32A KO 3.1, and ANP32A KO 3.2 cells collected at the indicated times. (**D**) Quantification of crystal-violet staining by absorbance at 590 nm. Data are mean ± SD from three independent experiments. (**E**) Representative propidium-iodide DNA-content profiles of WT and ANP32A-knockout cells. (**F**) Quantification of the percentages of cells in G0/G1, S, and G2/M phases and the apoptotic population. (**G**) Representative EdU/DNA-content flow-cytometry analysis of WT and ANP32A-knockout cells. (**H**) Quantification of cells in G0/G1, S, and G2 phases based on 15 min EdU incorporation and DNA content. Data in (**F**) and (**H**) are mean ± SD from three independent experiments. Data in (**D**), (**F**), and (**H**) were analyzed by two-way ANOVA followed by Dunn’s multiple-comparisons test. (**I** and **J**) Immunoblot analysis (**I**) and quantification (**J**) of cyclin D1 abundance relative to GAPDH. Data are mean ± SD from six independent biological replicates and were analyzed using Welch’s *t* tests. (**K-M**) Immunoblot analysis of RB and E2F1 (**K**) and quantification of E2F1 (**L**) and RB (**M**) relative to GAPDH. Data are mean ± SD from three independent biological replicates and were analyzed using two-tailed Welch’s t tests. Exact P values are indicated; ns, not significant.

**Figure 6:**
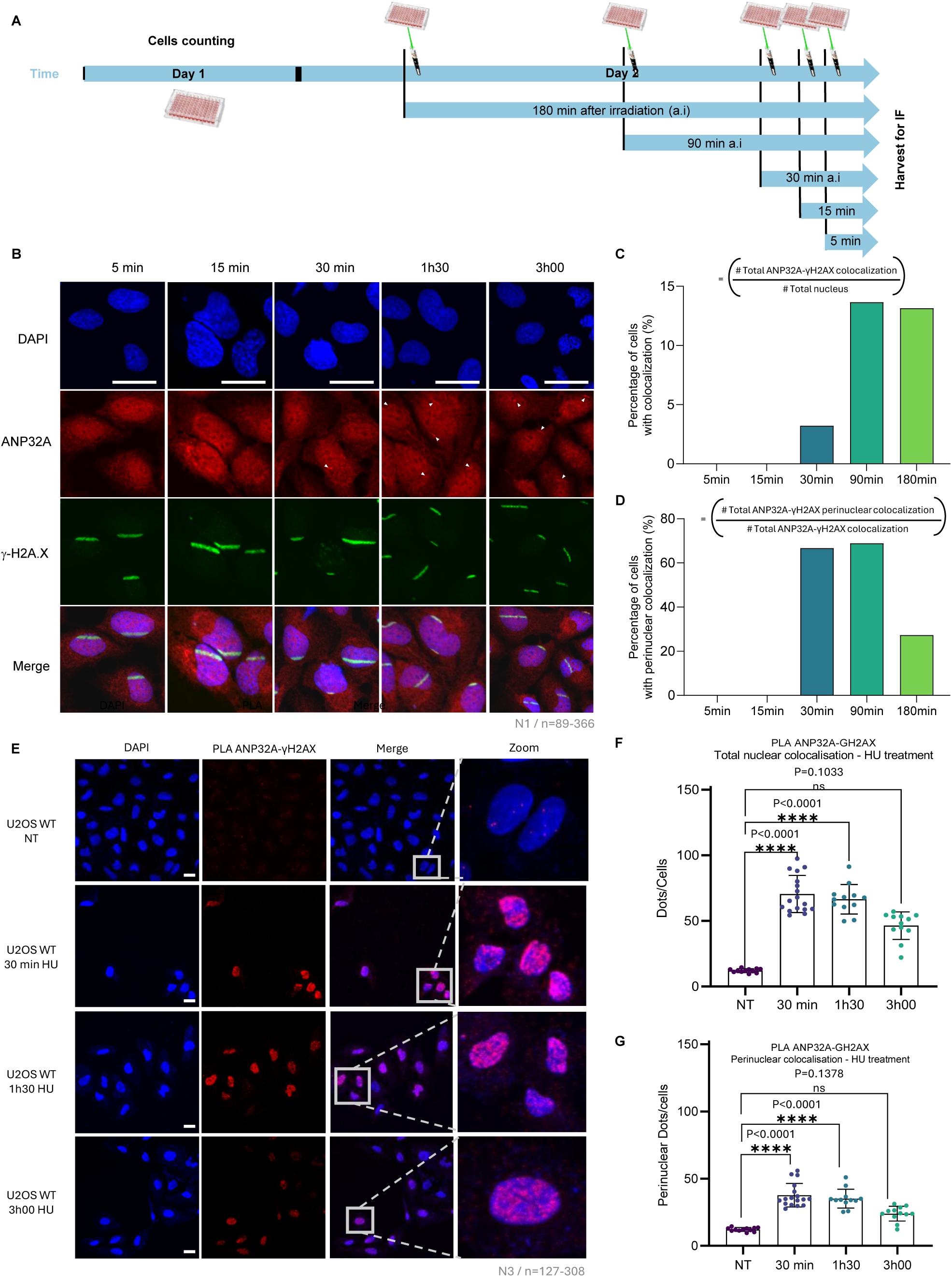
ANP32A is dynamically recruited to γH2AX-marked DNA lesions. (**A**) Experimental design for laser microirradiation. U2OS WT cells were subjected to localized laser-induced DNA damage and fixed 5 min, 15 min, 30 min, 90 min, or 180 min after irradiation. (**B**) Representative confocal images showing DNA stained with DAPI (blue), endogenous ANP32A (red), γH2AX (green), and merged channels at the indicated recovery times. Arrowheads indicate regions of ANP32A-γH2AX colocalization. Scale bars, 10 μm. (**C**) Percentage of cells displaying ANP32A-γH2AX colocalization at each recovery time. (**D**) Percentage of ANP32A-γH2AX colocalization events located at the nuclear periphery. The formulas used for quantification are shown in the figure. Data in (**C**) and (**D**) are from one independent experiment (N=1), with 89-366 cells analyzed per time point. (**E**) Representative PLA images showing endogenous ANP32A-γH2AX proximity in untreated U2OS WT cells and after 30 min, 90 min, or 180 min of HU treatment. PLA puncta are shown in red and nuclei in blue. Scale bars, 20 μm. (**F** and **G**) Quantification of ANP32A-γH2AX PLA puncta throughout the nucleus (**F**) and at the nuclear periphery (**G**). Data are mean ± SD from three independent biological replicates (N=3), with 127-308 cells analyzed per condition. Statistical significance was assessed using a Kruskal-Wallis test followed by Dunn’s multiple-comparisons test. Exact P values are indicated; ns, not significant.

To determine whether this proliferative defect was associated with altered cell-cycle progression, we analyzed DNA-content profiles by propidium iodide staining and flow cytometry. Compared with WT cells, both ANP32A-knockout clones displayed a marked increase in the G0/G1 population, accompanied by reduced representation of cells in S and G2/M phases (Figures 5E and 5F). Representative profiles from three independent experiments are shown in Figures S6A-S6C. We next used a 15-min EdU pulse combined with DNA-content analysis to identify actively replicating cells. Consistent with the DNA-content profiles, ANP32A-knockout cells exhibited an increased G0/G1 population and a reduced fraction of EdU-positive S-phase cells (Figures 5G, 5H and Figures S6D-S6F). Together, these complementary analyses indicate that ANP32A loss impairs G1/S progression and reduces the proportion of actively replicating cells.

To examine whether this phenotype was accompanied by changes in the G1/S regulatory machinery, we measured the abundance of cyclin D1, RB, and E2F1. Both ANP32A-knockout clones displayed increased cyclin D1 abundance relative to WT cells (Figures 5I and 5J), whereas RB and E2F1 levels were reduced (Figures 5K-5M). Given the known deregulation of the p16-RB-E2F axis in U2OS cells, these changes were not interpreted as evidence of direct regulation of the canonical RB-E2F pathway by ANP32A (Ma et al. 2003; Vernell et al. 2003). Rather, they indicate that ANP32A loss is associated with alterations in G1/S-associated regulatory proteins. Notably, the concomitant reduction in RB and E2F1 occurred despite the accumulation of ANP32A-deficient cells in G1 and their reduced progression into S phase, suggesting that these changes may reflect a secondary or compensatory response to impaired cell-cycle progression. These changes indicate that ANP32A loss disrupts the molecular network controlling the G1/S transition, although their causal contribution to the proliferation defect remains to be established. Collectively, these results show that ANP32A is required to sustain normal proliferation and cell-cycle progression.

### ANP32A is recruited to yH2AX-marked DNA lesions and enriched at the nuclear periphery

The enrichment of XRCC5 and XRCC6 in the replication stress-induced ANP32A proximal proteome, together with the increased proximity between ANP32A and XRCC5 following HU treatment, suggested that ANP32A may associate with damaged chromatin. To examine this possibility, we performed PLA between endogenous ANP32A and phosphorylated H2AX (γH2AX) in untreated U2OS cells and cells exposed to 1 mM HU for 24 h (Figure S7A). HU treatment increased the mean number of ANP32A-γH2AX PLA puncta from 51.6 to 100.0 per nucleus (Figure S7B). Signals localized at the nuclear periphery similarly increased from approximately 20.0 to 40.0 puncta per cell (Figure S7C). Thus, prolonged replication stress increases the proximity of ANP32A to γH2AX-marked chromatin.

To examine the spatial and temporal recruitment of ANP32A to localized DNA lesions, we next performed laser microirradiation followed by immunofluorescence analysis at 5 min, 15 min, 30 min, 90 min, and 180 min after irradiation (Figure 6A). ANP32A partially colocalized with γH2AX-marked damage tracks, with colocalization becoming apparent at 30 min, remaining prominent at 90 min, and decreasing by 180 min (Figures 6B and 6C). Among the ANP32A-γH2AX colocalization events detected at 30 min, 90 min, and 180 min, approximately 65%, 70%, and 25%, respectively, were located at the nuclear periphery (Figure 6D). These observations reveal temporally regulated recruitment of ANP32A to localized DNA lesions and an early enrichment of these events at the nuclear periphery.

We independently examined these recruitment kinetics by measuring ANP32A-γH2AX proximity following shorter periods of HU exposure (Figure 6E). Relative to 12.1 puncta per nucleus in untreated cells, PLA signals increased rapidly to 70.5 puncta after 30 min of HU treatment and subsequently declined to 66.5 and 46.4 puncta after 90 and 180 min, respectively (Figure 6F). Signals at the nuclear periphery followed a similar temporal profile, with the strongest enrichment observed after 30 min of HU exposure (Figure 6G). Together, the microirradiation and PLA analyses demonstrate that ANP32A is dynamically recruited to γH2AX-marked chromatin following DNA damage, with a substantial fraction of these associations occurring at the nuclear periphery. Combined with the HU-induced proximity of ANP32A to the NHEJ factors XRCC5 and XRCC6 identified by proximity proteomics, these findings further support a role for ANP32A in the DNA damage response and NHEJ-mediated repair.

### ANP32A promotes replication fork progression and NHEJ but is dispensable for stalled fork protection

Given that ANP32A loss impaired G1/S progression and that its proximal proteome included factors involved in DNA replication and repair, we asked whether ANP32A supports cell proliferation during replication-associated genotoxic stress. U2OS WT cells and the two ANP32A-knockout clones were exposed to HU, aphidicolin, etoposide, or cisplatin, and proliferation was monitored continuously for 120 h using the xCELLigence impedance system (Figure S8A). Under untreated conditions, ANP32A-knockout cells displayed lower cell indices at early time points, consistent with the growth defect detected by crystal violet staining (Figure S8B, 5C and 5D). Genotype-dependent differences were accentuated at selected time points during each treatment and were most pronounced and sustained following aphidicolin exposure (Figures S8C-S8F). These results indicate that ANP32A-deficient cells have reduced proliferative capacity during exposure to agents that perturb DNA replication and genome integrity.

To determine whether ANP32A directly affects DNA synthesis, we measured replication fork velocity by sequential CldU and IdU labeling under unperturbed conditions (Figure 7A). Replication forks progressed at an average velocity of 1.37 kb/min in WT cells, compared with 1.20 and 1.19 kb/min in ANP32A KO 3.1 and KO 3.2 cells, respectively (Figure 7B). Thus, ANP32A loss causes a moderate but significant reduction in replication-fork velocity, indicating that ANP32A supports efficient fork progression.

**Figure 7:**
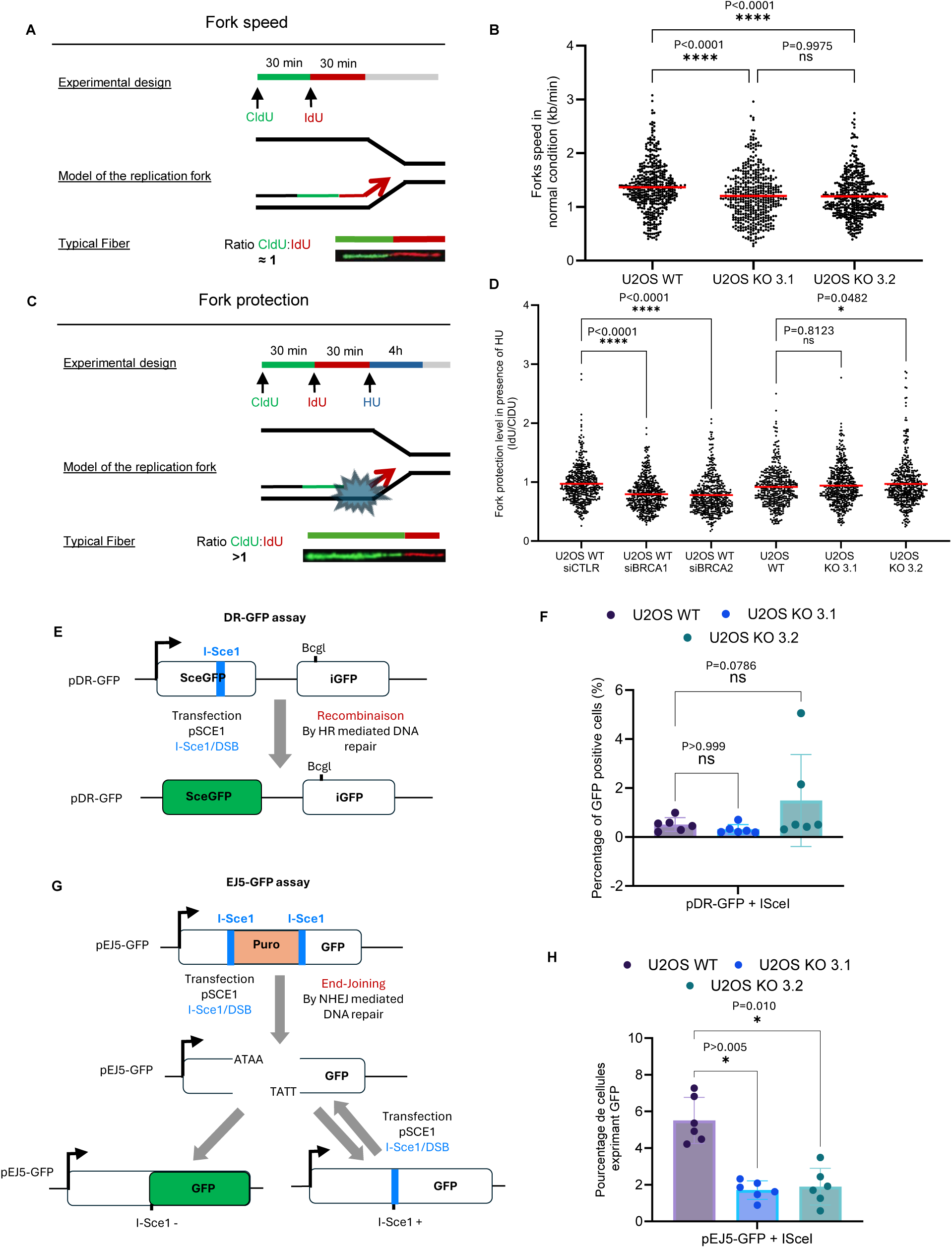
ANP32A loss slows replication forks and reduces end-joining reporter activity without impairing fork protection. (**A**) DNA-fiber assay used to measure replication-fork progression under unperturbed conditions. Cells were sequentially labeled with CldU (green; 100 μM, 30 min) and IdU (red; 250 μM, 30 min) before DNA-fiber spreading. A representative fiber containing consecutive CldU and IdU tracts is shown. (**B**) Replication fork velocities in U2OS WT, ANP32A KO 3.1, and ANP32A KO 3.2 cells. Each point represents an individual fiber, and horizontal red lines indicate mean fork velocity. At least 150 fibers were analyzed per biological replicate and condition. Pairwise comparisons were performed using two-way ANOVA with Šidák. (**C**) DNA-fiber assay used to measure stalled fork protection. Following sequential CldU and IdU labeling, cells were treated with 4 mM HU for 4 h. Fork protection was assessed using the IdU/CldU tract-length ratio; reduced ratios indicate degradation of newly synthesized IdU-labeled DNA. (**D**) IdU/CldU ratios in U2OS WT cells transfected with control, BRCA1, or BRCA2 siRNAs and in U2OS WT, ANP32A KO 3.1, and ANP32A KO 3.2 cells. BRCA1 and BRCA2-depleted cells served as positive controls for nascent-strand degradation. Each point represents an individual fiber, and horizontal red lines indicate the mean ratio. At least 150 fibers were analyzed per biological replicate and condition. (**E**) Schematic of the DR-GFP reporter used to measure homologous recombination. I-SceI cleavage of the inactive SceGFP sequence generates a double-strand break that can be repaired using the downstream iGFP sequence as a homologous template, restoring GFP expression. (**F**) DR-GFP reporter activity, measured as the percentage of GFP-positive cells after I-SceI expression. (**G**) Schematic of the EJ5-GFP reporter used to measure end joining. I-SceI cleavage at two sites excises the intervening puromycin resistance cassette, and end joining juxtaposes the promoter and GFP coding sequence to permit GFP expression. (**H**) EJ5-GFP reporter activity, measured as the percentage of GFP-positive cells after I-SceI expression. Data in (**F**) and (**H**) are mean ± SD from six independent biological replicates. Statistical significance was assessed using a Kruskal-Wallis test followed by Dunn’s multiple-comparisons test. Exact P values are indicated; ns, not significant.

We next distinguished this effect on ongoing fork progression from a potential role in protecting nascent DNA after fork stalling. Cells were sequentially labeled with CldU and IdU and then exposed to 4 mM HU for 4 h, after which fork protection was assessed using the IdU/CldU tract-length ratio (Figure 7C). As expected, depletion of BRCA1 or BRCA2 reduced the mean IdU/CldU ratio from 0.98 in siCTRL-treated cells to 0.79 and 0.78, respectively, validating the detection of nascent-strand degradation (Figure 7D). By contrast, the ratio was not reduced in either ANP32A-knockout clone: WT cells displayed a mean ratio of 0.92, compared with 0.94 and 0.97 in KO 3.1 and KO 3.2 cells, respectively. Immunoblotting confirmed ANP32A loss and efficient BRCA1 and BRCA2 depletion (Figures S9A and S9B). These findings demonstrate that ANP32A supports replication-fork progression but is dispensable for protecting nascent DNA following HU-induced fork stalling.

Because ANP32A was recruited to γH2AX-marked lesions and became proximal to the NHEJ factors XRCC5 and XRCC6 during replication stress, we next examined its contribution to double-strand break repair. Homologous recombination and NHEJ were measured using the DR-GFP and EJ5-GFP reporter systems, respectively (Figures 7E and 7G). ANP32A loss did not significantly alter HR reporter activity (Figure 7F) but reduced NHEJ reporter activity in both knockout clones (Figure 7H). Representative gating strategies from six biological replicates are shown in Figures S10A-S10F. These results identify a selective contribution of ANP32A to NHEJ and are consistent with the stress-induced proximity of ANP32A to XRCC5 and XRCC6.

## DISCUSSION

Here, we identify ANP32A as a chromatin-associated factor that supports two separable aspects of genome maintenance: efficient replication fork progression and non-homologous end joining. Time-resolved AirID proximity proteomics revealed phased remodeling of the ANP32A proximal proteome during HU-induced replication stress, with early enrichment of DNA replication and repair proteins followed by checkpoint and cell-cycle regulators. Orthogonal analyses revealed distinct stress-dependent proximity profiles between ANP32A and FEN1 or XRCC5, together with dynamic recruitment of ANP32A to γH2AX-marked DNA lesions. Functionally, ANP32A loss impaired G1/S progression, slowed replication forks, and reduced NHEJ reporter activity, while leaving stalled fork protection and homologous recombination reporter activity largely intact. These findings distinguish ANP32A from canonical fork protection factors and position it at the interface between chromatin organization, DNA replication, and double-strand break repair.

Time-resolved proximity mapping further showed that ANP32A does not occupy a static molecular environment during replication stress. Early HU exposure enriched ANP32A-proximal proteins involved in DNA replication and repair, including FEN1, MCM2, PRIM1, RNASEH2B, XRCC5, and XRCC6, whereas prolonged stress shifted the network toward checkpoint, cell-cycle, and mitotic regulators. This phased remodeling is consistent with previous iPOND and proximity-proteomic studies showing extensive temporal reorganization of stalled replisomes (Dungrawala et al. 2015; Jurkovic et al. 2024). The distinct PLA kinetics of FEN1 and XRCC5 further suggest that ANP32A associates with different DNA-metabolic environments as replication stress evolves. Sustained proximity to FEN1 is consistent with its functions in Okazaki fragment maturation and the processing and restart of stressed replication forks, whereas the transient XRCC5 response may reflect early engagement with Ku-containing complexes at arrested or broken replication intermediates (Balestrini et al. 2013; Teixeira-Silva et al. 2017; Xu et al. 2018). Because prolonged HU exposure can convert a subset of stalled forks into damaged or collapsed structures, these temporal changes may reflect redistribution of ANP32A between replication-associated and repair-associated chromatin rather than assembly of a single stable complex (Petermann et al. 2010). Moreover, because AirID and PLA measure proximity rather than direct binding, determining whether ANP32A recruits, stabilizes, or regulates these factors at specific DNA structures will require biochemical and fork-resolved analyses.

The cellular phenotypes of ANP32A loss further support a role in DNA replication, while suggesting contributions to both cell-cycle progression and DNA synthesis. ANP32A-deficient cells accumulated in G0/G1, contained fewer EdU-positive S-phase cells, and displayed increased cyclin D1 abundance together with reduced RB and E2F1. Because these changes do not conform to a simple linear inhibition of the cyclin D-RB-E2F axis, they may reflect compensatory remodeling of G1/S regulatory pathways rather than the direct cause of the cell-cycle defect. Previous work showed that depletion of PP32/ANP32A disrupts the maturation of newly synthesized histone H4 by permitting premature HAT1-dependent acetylation, thereby destabilizing H4 and causing S-phase accumulation (Saavedra et al. 2017). Although the precise cell-cycle distribution differs between the two studies, both findings connect ANP32A-dependent histone regulation to efficient S-phase progression. Our DNA fiber analyses extend this connection by showing that ANP32A loss also reduces the velocity of individual replication forks. Because replication fork speed depends on an adequate supply of newly synthesized histones and their efficient assembly into chromatin, defective H4 maturation provides a plausible mechanism linking ANP32A loss to slower DNA synthesis (Mejlvang et al. 2014). This possibility remains to be tested directly, however, as histone deposition and nucleosome assembly were not measured in our system. Importantly, ANP32A loss did not promote nascent strand degradation following HU-induced fork stalling, distinguishing its role in ongoing fork progression from the fork protection functions of factors such as BRCA2 (Schlacher et al. 2011). Together, these observations suggest that ANP32A facilitates DNA replication potentially by supporting a chromatin environment compatible with efficient fork movement rather than by directly protecting stalled forks.

The recruitment of ANP32A to γH2AX-marked lesions and the reduction in end-joining activity following ANP32A loss connect its stress-regulated proximal network to a functional DNA repair outcome. Both HU treatment and laser microirradiation induced rapid but transient proximity or colocalization between ANP32A and γH2AX, consistent with the redistribution of ANP32A to damaged chromatin. The stress-dependent enrichment of XRCC5 and XRCC6, together with increased ANP32A-XRCC5 proximity, further points to a Ku-associated repair process. Ku70-Ku80 is a central DNA-end recognition complex in classical NHEJ but also contributes to the processing of replication-associated single-ended breaks and arrested replication forks (Balestrini et al., 2013; Teixeira-Silva et al., 2017). Accordingly, the reduced EJ5-GFP activity observed in ANP32A-deficient cells, despite preserved DR-GFP activity, suggests a selective contribution to end joining rather than a general defect in double-strand break repair. However, because EJ5-GFP detects multiple classes of end-joining events, these experiments do not establish whether ANP32A acts specifically in classical NHEJ or at the level of DNA end recognition, processing, synapsis, or ligation (Bennardo et al. 2008). Given the established chromatin regulatory functions of ANP32A, one possibility is that it creates or maintains a chromatin environment that facilitates efficient Ku-dependent repair. The enrichment of ANP32A-γH2AX signals at the nuclear periphery is also intriguing because nuclear positioning and local chromatin state can influence repair-pathway use (Lemaître et al. 2014; Schep et al. 2021; Nesic et al. 2025). Nevertheless, determining whether these lesions are associated with heterochromatin or lamina-associated domains will require direct analysis using markers such as H3K9me3, HP1α, or lamins.

Together, our findings support a model in which ANP32A helps maintain a chromatin environment conducive to efficient DNA replication and repair. Under unperturbed conditions, ANP32A promotes G1/S progression and replication fork movement, potentially through its established role in histone homeostasis, whereas replication stress remodels its proximal network to include factors involved in DNA processing and Ku-dependent end joining. These activities appear functionally distinct: ANP32A loss slowed ongoing forks without compromising stalled fork protection and reduced end joining without measurably affecting homologous recombination. Several questions remain, including whether ANP32A directly associates with FEN1 or Ku, which ANP32A domains mediate these proximity relationships, and whether altered chromatin assembly provides a common mechanistic basis for the replication and repair phenotypes. Addressing these questions will require rescue experiments with separation-of-function mutants, biochemical analysis of candidate complexes, and validation in additional cellular and physiological models. Nevertheless, by combining time-resolved proximity proteomics with functional analyses, this study defines a previously unrecognized contribution of ANP32A to replication fork progression and NHEJ and expands the role of the ANP32 protein family in mammalian genome maintenance.

## Acknowledgments

We thank Véronique Giroux, Maryline Labrie and Marie-Josée Bouchée for providing antibodies used in this study. We also thank Daniel Garneau for assistance with confocal and fluorescence microscopy and automated image analysis; Mathieu Catala for flow cytometry training and assay optimization; and Sarah Dubois, from the laboratory of Steve Jean, for assistance in developing the CellProfiler analysis pipeline.

## Author contributions

Conceptualization, C.-M.J. and F.-M.B.; Methodology, C.-M.J., J.R., G.M., D.L., I.M., B.D. and A.M.; Investigation, C.-M.J., J.R., G.M., D.L., I.M. and B.D.; Formal analysis, C.-M.J., J.R., G.M., D.L., I.M., B.D. and A.D.D.M.; Data curation, C.-M.J. and D.L.; Visualization, C.-M.J.; Resources, A.M. and F.-M.B.; Writing - original draft, C.-M.J. and F.-M.B.; Writing - review & editing, C.-M.J., J.R., D.L., I.M., A.M., and F.-M.B.; Supervision, A.M. and F.-M.B.; Project administration, C.-M.J. and F.-M.B.; Funding acquisition, F.-M.B.

## Funding and additional information

This work was supported by the Natural Sciences and Engineering Research Council of Canada (NSERC; RGPIN-2024-05256) to F.-M.B. F.-M.B. is supported by a Distinguished Research Scholar Award from the Fonds de recherche du Québec - Santé (FRQS; #366044). C.-M.J. is the recipient of a doctoral scholarship from FRQS and a Doctoral Research Award from the Cancer Research Society.

---

**Supplementary Figure S1:**
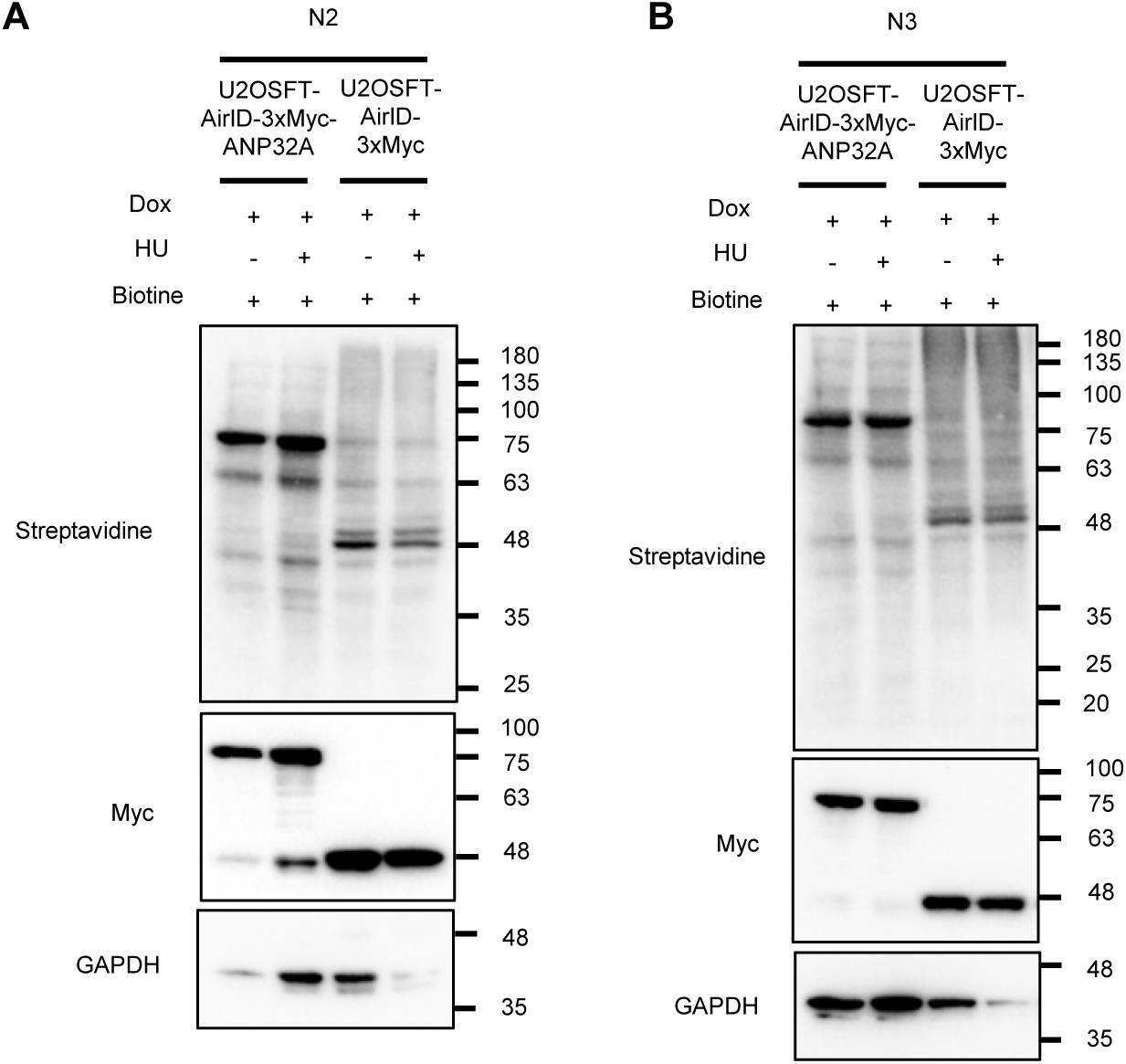
Independent validation of the AirID proximity-labeling system. (**A** and **B**) Immunoblot validation of AirID-3×Myc-ANP32A and AirID-3×Myc expression and proximity biotinylation in independent biological replicates N2 (**A**) and N3 (**B**). Cells were induced with doxycycline and supplemented with biotin in the absence or presence of 1 mM HU for 24 h. Biotinylated proteins were detected using streptavidin-HRP, AirID constructs using an anti-Myc antibody, and GAPDH as a loading control. Related to Figure 1.

**Supplementary Figure S2:**
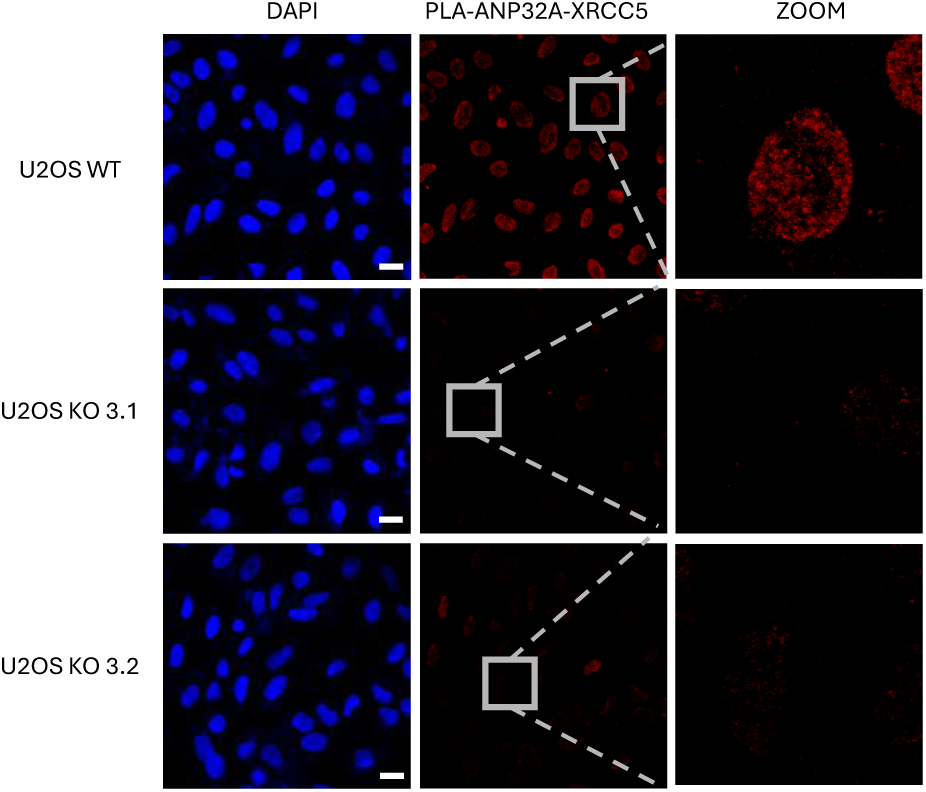
Genetic validation of ANP32A-XRCC5 PLA specificity. Representative confocal images showing endogenous ANP32A-XRCC5 PLA in U2OS WT cells and two independently isolated ANP32A-knockout clones, KO 3.1 and KO 3.2. PLA puncta are shown in red and nuclei in blue. Enlarged views of representative nuclei are shown on the right. Scale bar, 20 μm. Related to Figure 2.

**Supplementary Figure S3:**
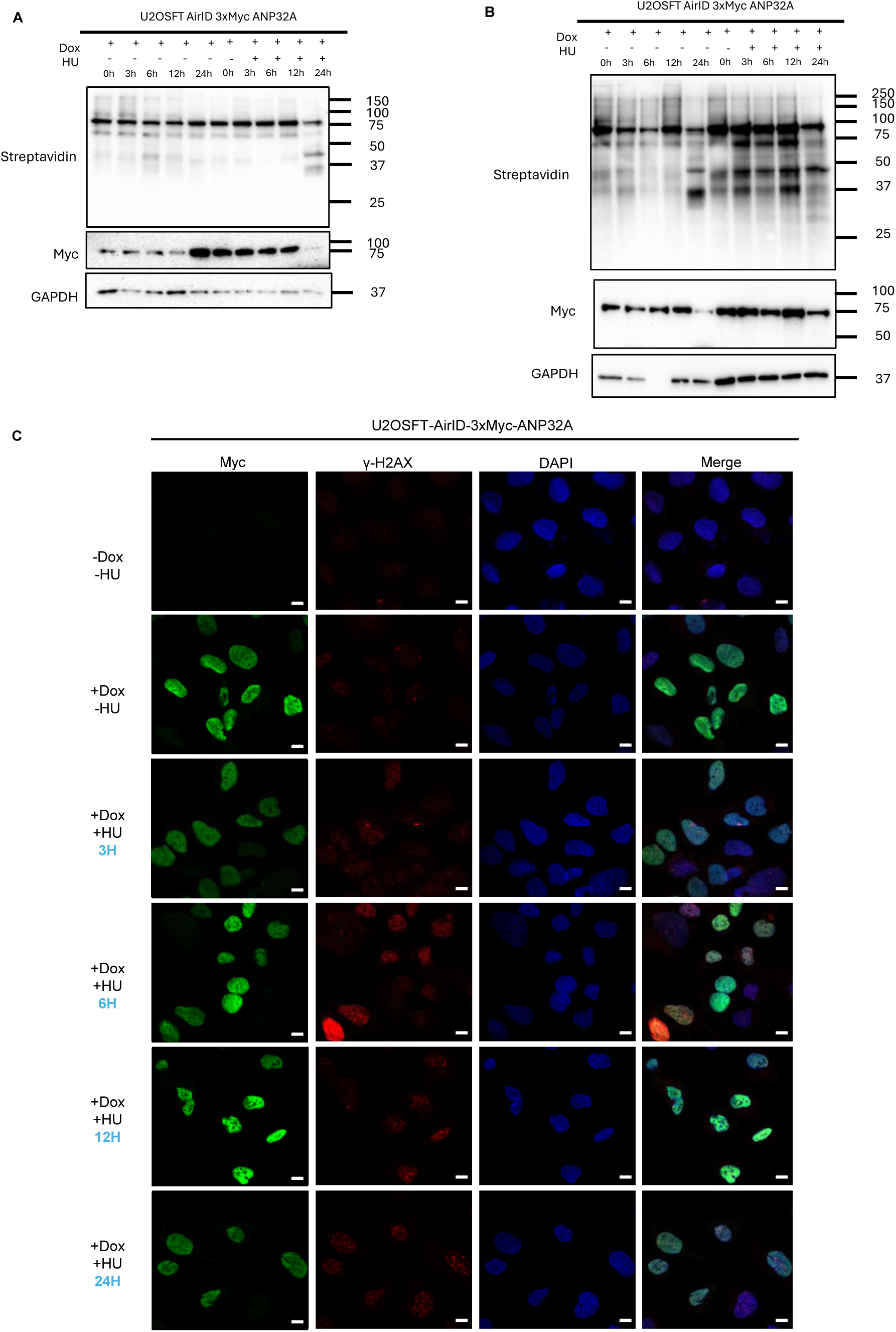
Validation of time-resolved AirID labeling and HU-induced replication stress. (**A** and **B**) Immunoblot analysis of AirID-3×Myc-ANP32A expression and proximity biotinylation in matched untreated and HU-treated samples collected at the indicated time points. Biotinylated proteins were detected using streptavidin-HRP, the fusion protein using an anti-Myc antibody, and GAPDH as a loading control. Panels show independent biological replicates N2 (**A**) and N3 (**B**). (**C**) Immunofluorescence analysis of AirID-3×Myc-ANP32A expression and localization and γH2AX accumulation under the indicated doxycycline and HU-treatment conditions. Myc is shown in green, γH2AX in red, and DNA in blue. Scale bars, 5 μm. Related to Figure 3.

**Supplementary Figure S4:**
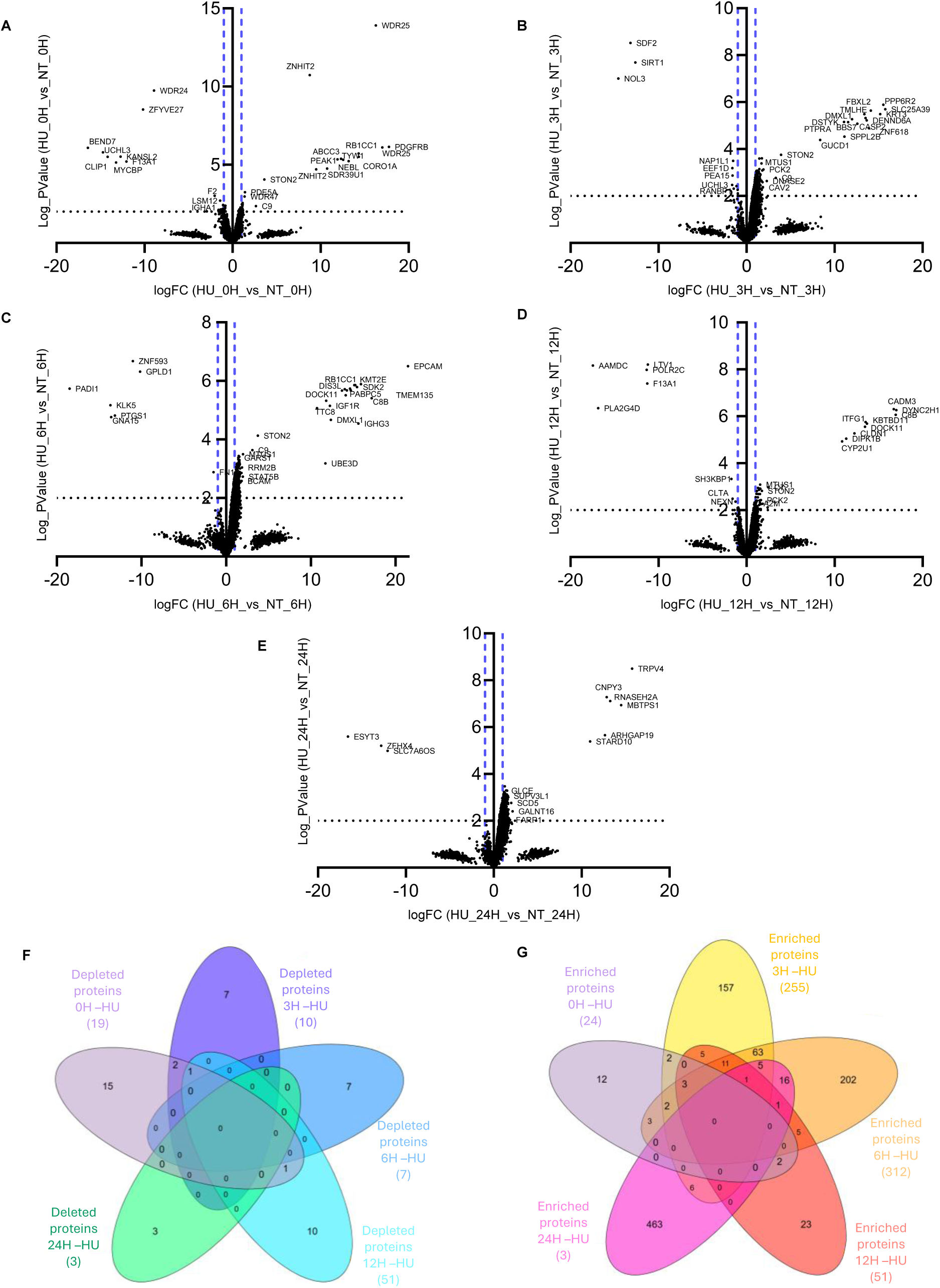
Differential analysis of time-resolved ANP32A proximal proteome. (**A-E**) Volcano plots comparing ANP32A-proximal protein abundance between HU-treated and matched untreated samples collected at 0 h (**A**), 3 h (**B**), 6 h (**C**), 12 h (**D**), or 24 h (**E**). The x axis represents log2 fold change between HU-treated and untreated samples, and the y axis represents -log10(P value). Dashed vertical lines denote log2 fold-change thresholds of -1 and +1, and the dotted horizontal line denotes the statistical threshold corresponding to an FDR ≤5%. (**F** and **G**) Venn diagrams showing the overlap among significantly depleted (**F**) and enriched (**G**) proteins at the different time points. Related to Figure 3.

**Supplementary Figure S5:**
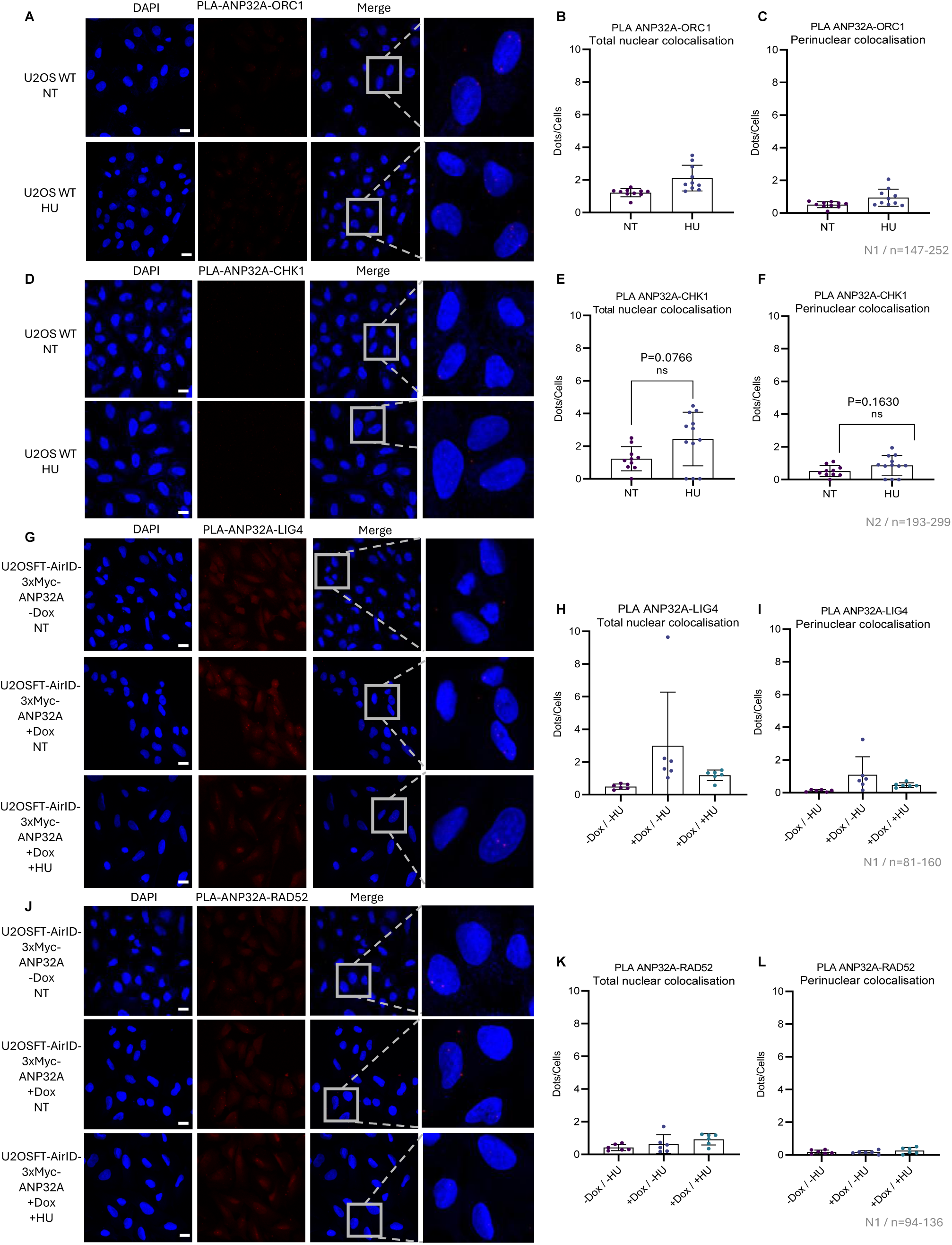
PLA analysis of proteins not enriched in the ANP32A proximal proteome. ORC1, CHK1, LIG4, and RAD52 did not meet the enrichment criteria in the time-resolved AirID analysis. (**A**-**C**) Representative images of endogenous ANP32A-ORC1 PLA in untreated U2OS WT cells and following 24 h of HU treatment (**A**), with quantification of puncta throughout the nucleus (**B**) and at the nuclear periphery (**C**). (**D-F**) Representative images of endogenous ANP32A-CHK1 PLA under the same conditions (**D**), with quantification of total nuclear (**E**) and perinuclear (**F**) puncta. (**G-I**) Representative ANP32A-LIG4 PLA images in U2OSFT AirID-3×Myc-ANP32A cells cultured without doxycycline, with doxycycline, or with doxycycline and HU (**G**), with quantification of total nuclear (**H**) and perinuclear (**I**) puncta. **(J-L)**. Representative ANP32A-RAD52 PLA images and corresponding quantification under the same conditions. PLA puncta are shown in red and nuclei in blue. Each point represents one analyzed image, and data are mean ± SD. The numbers of independent experiments and analyzed cells are indicated in the figure. For CHK1, comparisons between untreated and HU-treated conditions were performed using two-tailed Mann-Whitney tests; ns, not significant. Scale bars, 20 μm. Related to Figure 4.

**Supplementary Figure S6:**
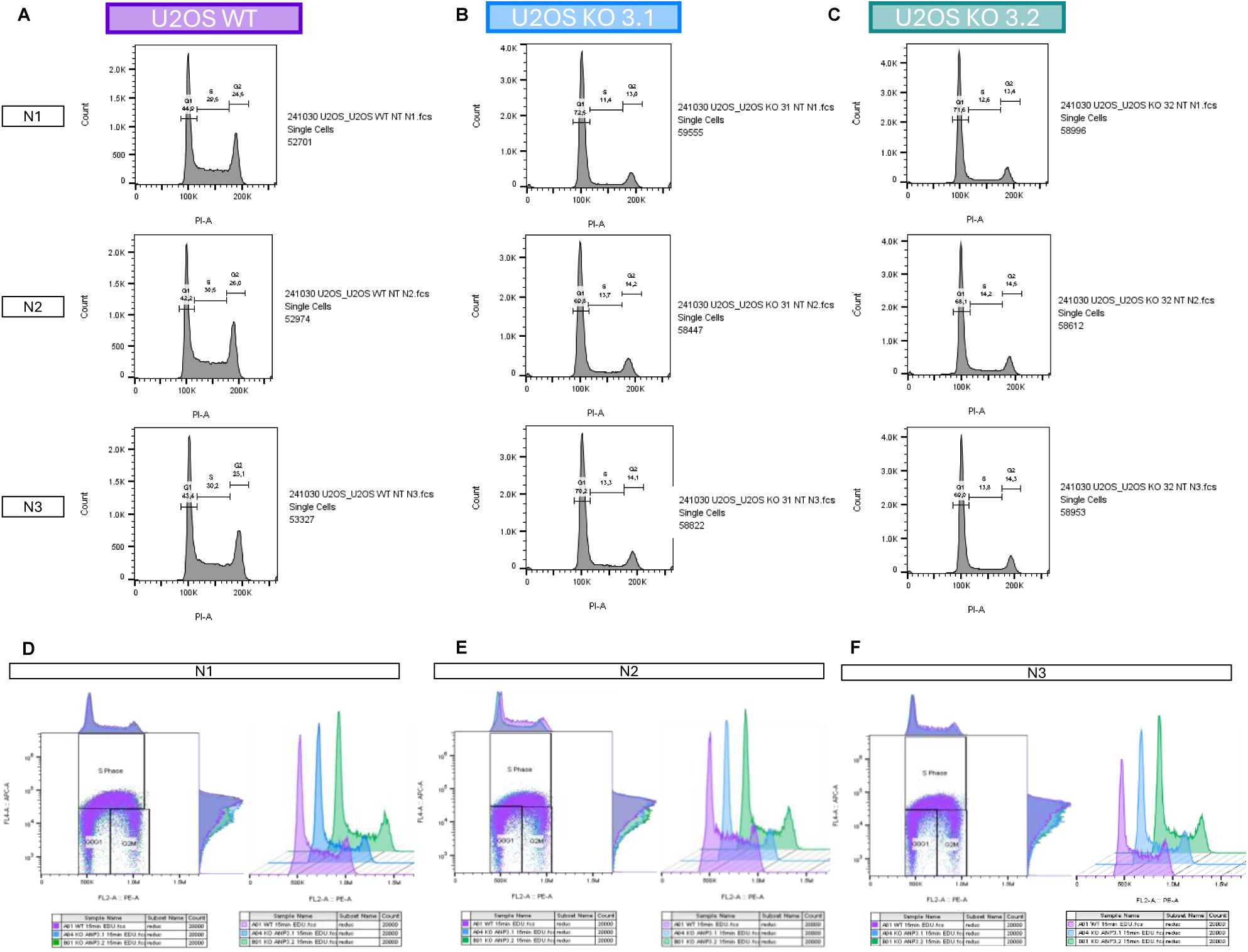
Cell-cycle profiles of U2OS WT and ANP32A-knockout cells. (**A-C**) Propidium-iodide DNA-content histograms and cell-cycle modeling for U2OS WT (A), ANP32A KO 3.1 (**B**), and ANP32A KO 3.2 (**C**) cells. Each panel shows the three independent biological replicates, N1-N3, used for the quantification in Figure 5F. (**D-F**) EdU incorporation and DNA-content profiles from independent biological replicates N1 (**D**), N2 (**E**), and N3 (**F**). Each panel shows the profiles of U2OS WT, ANP32A KO 3.1, and ANP32A KO 3.2 cells used for the quantification in Figure 5H. Related to Figure 5.

**Supplementary Figure S7:**
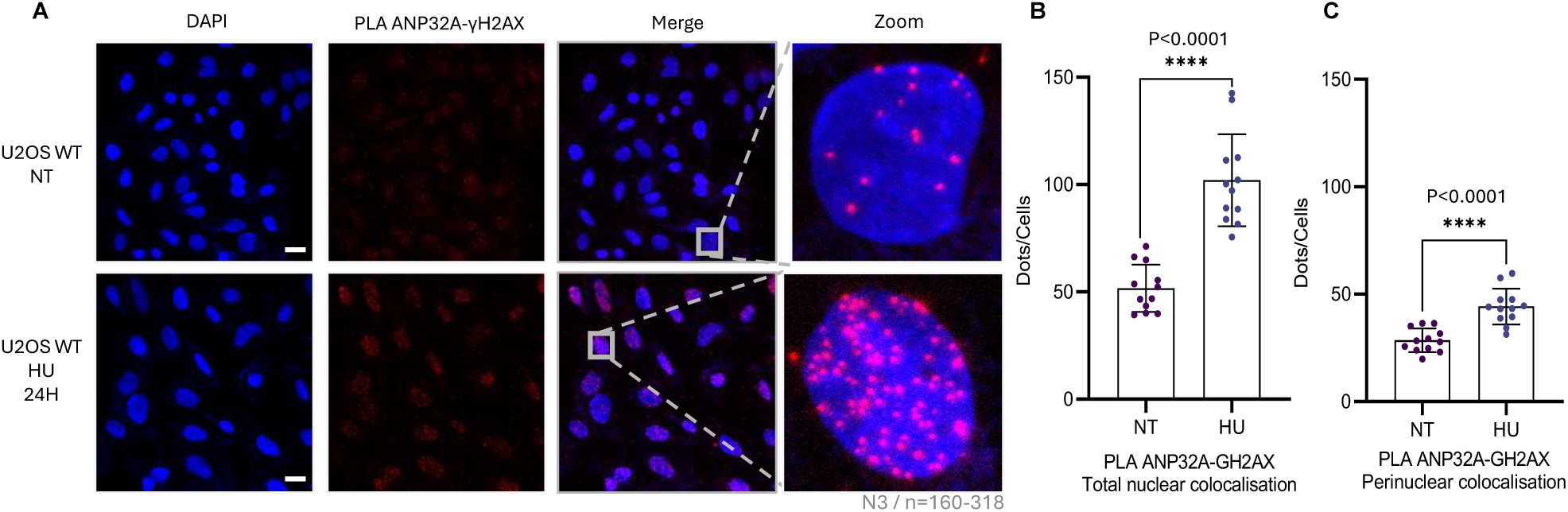
Replication stress increases ANP32A proximity to γH2AX-marked chromatin. (**A**) Representative PLA images showing endogenous ANP32A-γH2AX proximity in untreated U2OS WT cells and following treatment with 1 mM HU for 24 h. PLA puncta are shown in red and nuclei in blue. (**B** and **C**) Quantification of ANP32A-γH2AX PLA puncta throughout the nucleus (**B**) and at the nuclear periphery (**C**). Data are mean ± SD from three independent biological replicates (N=3), with 160-318 cells analyzed per condition. Statistical significance was assessed using two-tailed Mann-Whitney tests. Scale bars, 20 μm. Related to Figure 6.

**Supplementary Figure S8:**
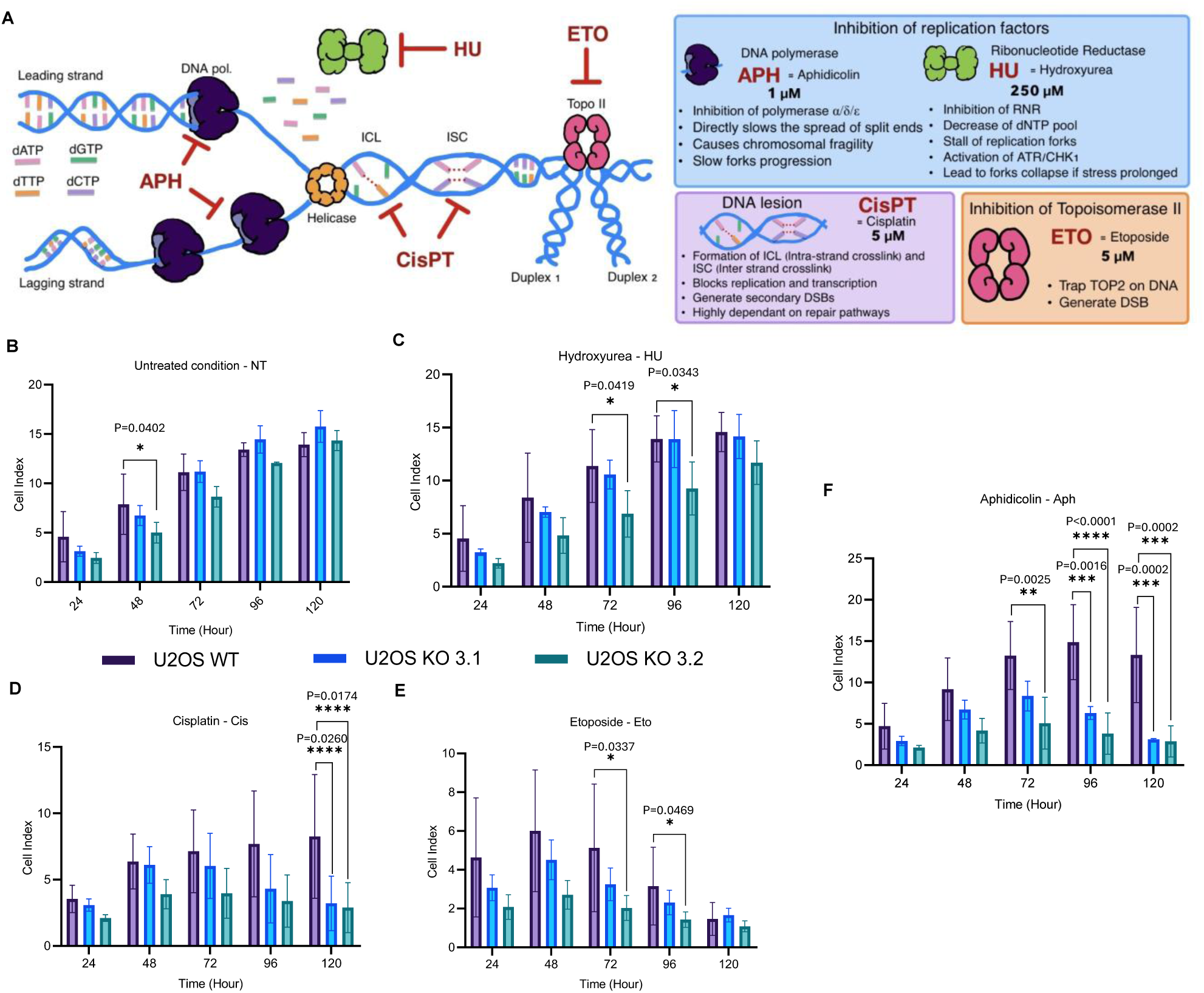
Proliferation of ANP32A-deficient cells during genotoxic stress. (**A**) Schematic showing the principal effects of the agents used to perturb DNA replication and genome integrity. (**B-F**) Real-time proliferation of U2OS WT, ANP32A KO 3.1, and ANP32A KO 3.2 cells measured using the xCELLigence impedance system under untreated conditions (**B**) or during exposure to 250 μM HU (**C**), 5 μM cisplatin (**D**), 5 μM etoposide (**E**), or 1 μM aphidicolin (**F**). Treatments were initiated 24 h after seeding, and cell index was monitored for 120 h. Data are mean ± SD from three independent biological replicates. Statistical significance was assessed using two-way ANOVA followed by Dunnett’s multiple-comparisons test. Only significant comparisons and their exact P values are indicated. Related to Figure 7.

**Supplementary Figure S9:**
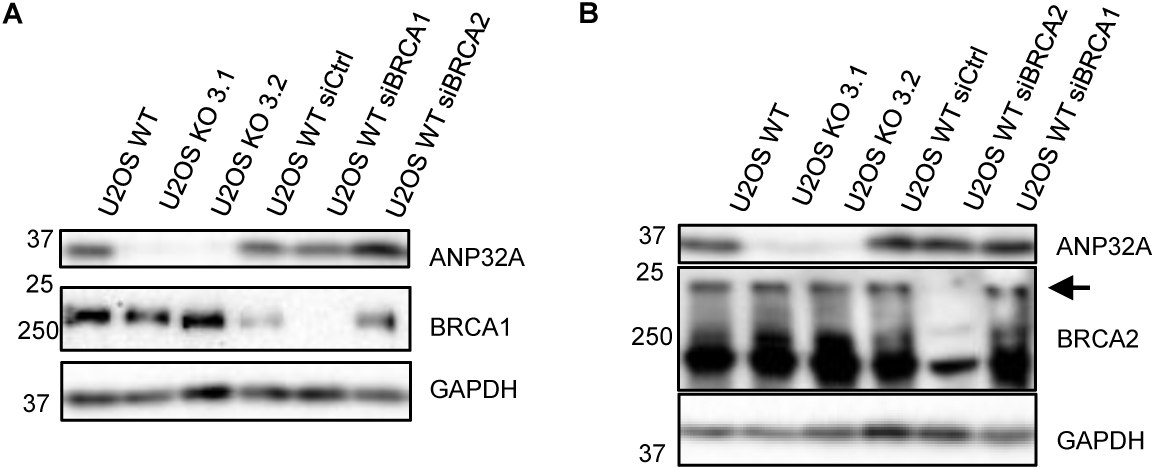
Validation of ANP32A genotype and BRCA1/BRCA2 depletion in the fork-protection assay. (**A** and **B**) Immunoblot analysis of U2OS WT, ANP32A KO 3.1, and ANP32A KO 3.2 cells and U2OS WT cells transfected with control, BRCA1, or BRCA2 siRNAs. ANP32A and BRCA1 expression are shown in (**A**), and ANP32A and BRCA2 expression in (**B**). GAPDH was used as a loading control. The arrow indicates the BRCA2 band. Related to Figure 7.

**Supplementary Figure S10:**
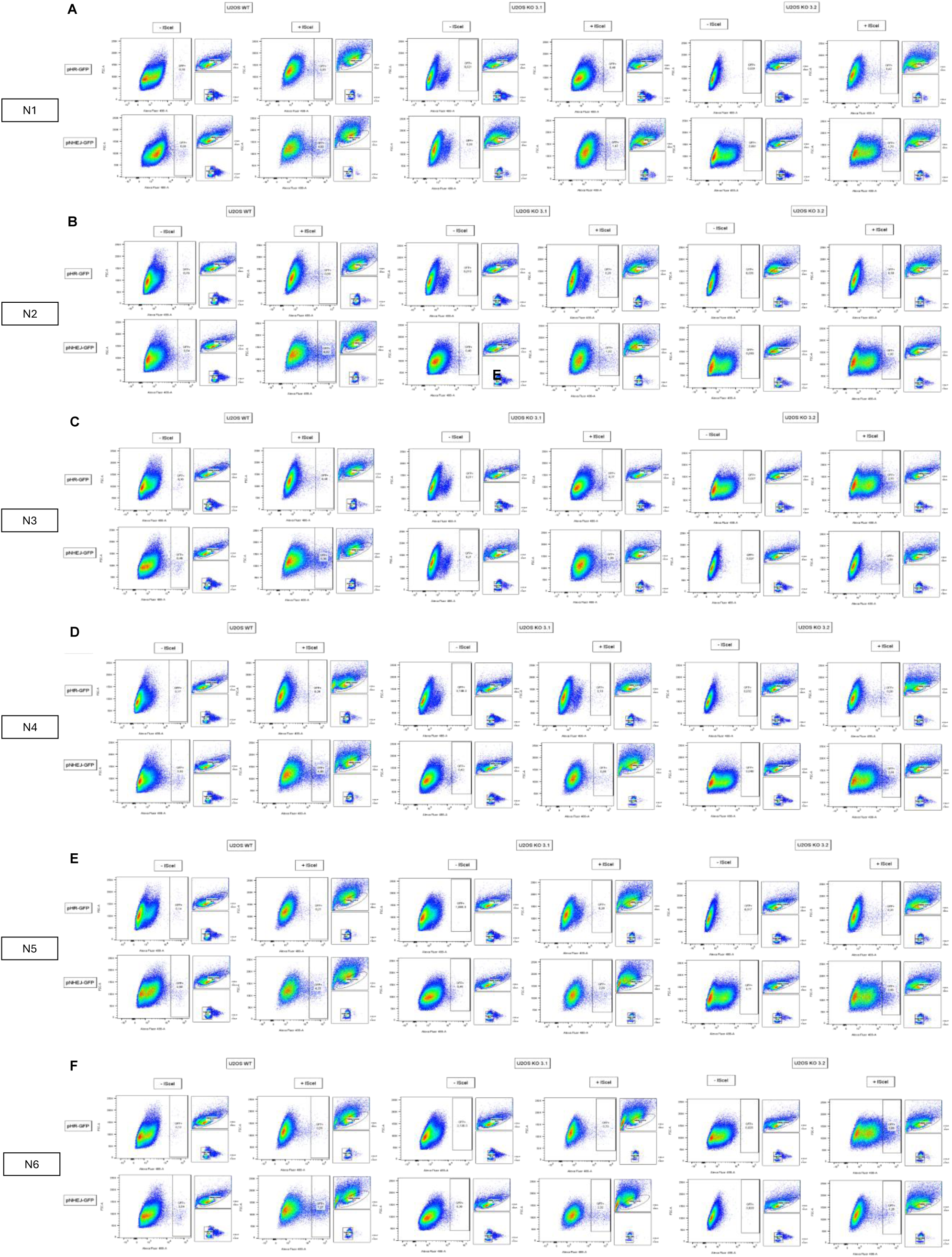
Flow-cytometry gating for the DR-GFP and EJ5-GFP reporter assays. (**A-F**) Flow cytometry gating applied to six independent biological replicates, N1-N6, of U2OS WT, ANP32A KO 3.1, and ANP32A KO 3.2 cells stably carrying the DR-GFP or EJ5-GFP reporter. Each genotype is shown with and without transient I-SceI expression. Sequential gates based on forward and side scatter were used to exclude debris, select singlets, and identify GFP-positive cells. The resulting percentages of GFP-positive cells were used to quantify DR-GFP and EJ5-GFP reporter activity in Figures 7F and **7H**, respectively.

